# Evolution and antiviral activity of a human protein of retroviral origin

**DOI:** 10.1101/2020.08.23.263665

**Authors:** John A. Frank, Manvendra Singh, Harrison B. Cullen, Raphael A. Kirou, Meriem Benkaddour-Boumzaouad, Jose L. Cortes, Jose Garcia-Perez, Carolyn B. Coyne, Cédric Feschotte

## Abstract

Endogenous retroviruses are abundant components of mammalian genomes descended from ancient germline infections. In several mammals, the envelope proteins encoded by these elements protect against exogenous viruses, but this activity has not been documented in human. We report that our genome harbors a large pool of envelope-derived sequences with the potential to restrict retroviral infection. To further test this, we characterize in detail the envelope-derived protein, *Suppressyn*. We found that *Suppressyn* is expressed in preimplantation embryos and developing placenta using its ancestral retroviral promoter. Restriction assays in cell culture show that *Suppressyn*, and its hominoid orthologs, can restrict infection by extant mammalian type D retroviruses. Our data support a generalizable model of retroviral envelope cooption for host immunity and genome defense.

**Summary:** We found that the human genome expresses a vast pool of envelope sequences of retroviral origin and provide proof of principle that such proteins can restrict zoonotic viruses.

---

Viruses pose a persistent threat to human health and may be potent evolutionary drivers of immune adaptation (*1*–*3*). Thus, it is critical to identify host factors that protect against viral infection. Endogenous retroviruses (ERV) are abundant components of mammalian genomes that descend from ancient retroviral germline infections (*1*). ERV sequences are a source of novel genes in evolution (*2*). ERV-derived envelopes (*env*) are particularly interesting because they can serve as antiviral factors by binding to and competing for target cell receptors of exogenous viruses (*1*). Env-derived proteins restricting against contemporary infectious retroviruses have been documented in mouse (Fv4), cat (Refrex-1), and sheep (enJSS6A1) but not in humans (*1, 2, 4*). Importantly, the receptor-binding surface domain of an *env* can be sufficient to restrict infection by this interference mechanism (*4*). Because the human genome contains thousands of remnant retroviral sequences with coding potential, we hypothesized that it includes envelope sequences with the capacity to restrict against modern viruses.

To assess the pool of potentially antiviral sequences, we scanned the human genome for *env*-derived open reading frames (*envORF)* that encode at least 70 aa and are predicted to include the receptor-binding surface domain (see Methods). This search identified a total of 1,507 unique *envORFs*, including ∼20 *env*-derived sequences currently annotated as human genes such as *Supppressyn* (*SUPYN*), *Syncytin-1* and *Syncytin-2* (*5, 6*). This number is consistent with a previous genome-wide scan for ORFs derived from endogenous viral elements predicting 1,731 ORFs with homology to retroviral envelopes (*7*). Next, we mined transcriptome datasets generated from human embryos and various adult tissues **(Table S1)** and observed that ∼44% of *envORF*s (668/1507) showed evidence of RNA expression in at least one of the cell types surveyed **(Fig 1a)**. These analyses revealed three general trends about expressed *envORFs*: (i) like known *env-*derived genes, *envORFs* exhibit tissue-specific expression patterns **(Fig 1a; fig S1, S2)**; (ii) the majority of *envORFs* are expressed during early human development, often in stem and progenitor germ cells, and/or in placenta **(Fig 1a; fig S1)**; (iii) with the exception of brain, *envORFs* are rarely expressed in differentiated tissues under normal conditions **(Fig 1a; fig S1, S2)**. Since antiviral factors are generally expressed in immune cells and/or induced upon immune stimulation or infection, we also profiled *envORF* expression using transcriptome datasets generated from a variety of immune cells, including resting, stimulated, and HIV-infected cells **(Table S1)**. These analyses revealed that a substantial fraction of *envORFs* (145 loci from multiple families) are expressed in immune cells and tend to be induced by immune simulation **(Fig 1b; fig S3)**. Consistent with previous reports (*8*), we also observed that a subset of *envORFs* are induced upon HIV infection **(Fig 1b, c; fig. S3, S4)**. Together, these data suggest that the human genome harbors a vast reservoir of *env*-derived sequences with coding potential for receptor-binding proteins, many of which are expressed in a tissue-specific or infection-inducible fashion, suggesting that they may function as antiviral factors.

**Figure 1:**
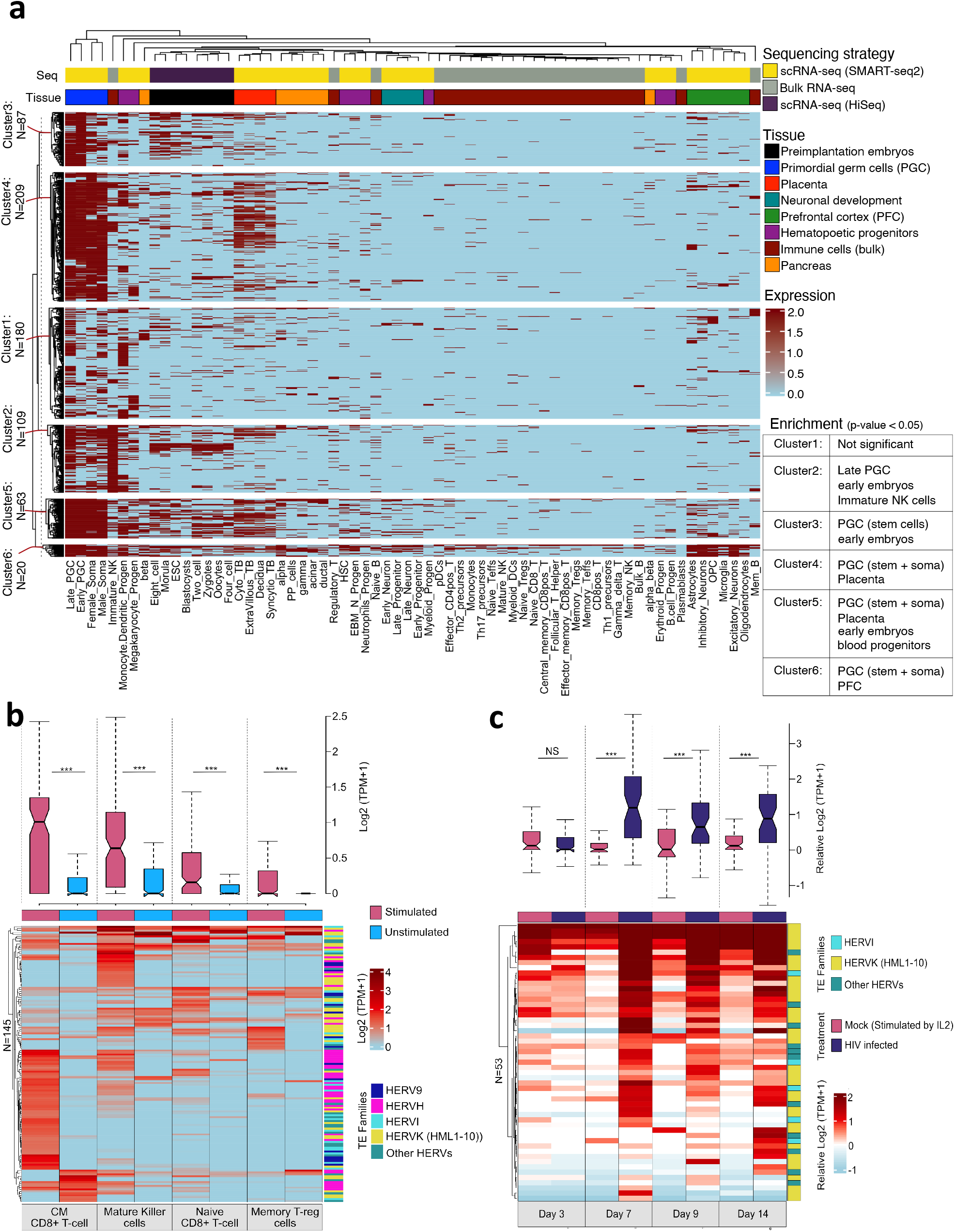
Expression profile of env-derived transcripts over a subset of human cell types. **(a-c)** Heatmaps show *envORF* expression in 66 distinct cell-types from 8 independent datasets (a), and in response to stimulation (b) and HIV infection in immune cells (c). Rows and columns represent individual *envORF* loci and cell identity respectively. *EnvORF* loci were included in heatmap if log2 CPM > 1 in at least one cell type. Bars located above heatmaps denote sequencing strategy (a, top), tissue source (a: bottom) or treatment (b, c) with the same color scheme shown in figures S1, S3, and S4 respectively. Rows and columns are ordered by hierarchical clustering based on *envORF* expression. Significant *envORF* enrichment in indicated clusters was calculated by hypergeometric test. **(b, c)** Boxplots represent the distribution of envORF expression levels relative to unstimulated (b) and Mock-infected cells (c) (NS: not significant; ***p < 0.01; Wilcoxon rank-sum test)

As a paradigm to test the antiviral activity of an envORF, we focused on SUPYN for two primary reasons. First, SUPYN was reported to be expressed in the developing placenta, as we observe for many envORFs, a tissue that can be vulnerable to viral infection and a barrier against vertical transmission to the fetus (*9*). Second, *SUPYN* lacks a transmembrane domain but retains the ability to bind the amino acid transporter ASCT2 (also known as SLC1A5), which is the receptor for a diverse group of retroviruses, including endogenous (e.g. HERV-W) and exogenous viruses such as RD114 in cat and Simian Retrovirus (SRV) in Old World monkeys (*5, 10*–*12*). These viruses are collectively referred to as RD114 and Type D (RDR) retroviruses (*12*). Thus, we hypothesized that SUPYN might restrict RDR retroviruses by virtue of its binding to ASCT2.

To obtain a more detailed view of *SUPYN* expression and regulation in adult tissues and during human development, we analyzed publicly available bulk and single-cell RNA-seq, ATAC-seq, DNase-seq, and ChIP-seq datasets generated from adult tissues, preimplantation embryos, human embryonic stem cell lines (hESC), *in vitro* trophoblast (TB) differentiation models, and placenta explants isolated at multiple stages of pregnancy **(Table S1)**. Apart from testis and cerebellum, *SUPYN* expression is generally low (<1 log2 (TPM+1)) or absent in adult tissues (**fig S2**). Conversely, *SUPYN* is expressed at high levels in preimplantation embryos from the 8-cell stage of development **(Fig 2a, fig S5a)**. Accordingly, the *SUPYN* promoter region, including the long terminal repeat (LTR) of the ancestral HERVH48 provirus, is marked by several peaks of accessible chromatin from the 8-cell to blastocyst stages **(fig S5b)**. In hESCs, *SUPYN* RNA is also abundant (fig 2a) and its promoter region is marked by histone modifications characteristic of transcriptionally active chromatin (H3K4me1, H3K27ac) and bound by core pluripotency (OCT4, NANOG, KLF4, SMAD1) and self-renewal (SRF, OTX2) transcription factors **(Fig 2b)**. During hESC to TB differentiation, pluripotency factors (NANOG, OCT4) occupying the *SUPYN* promoter region are replaced by TB-specific transcription factors (TFAP2A, GATA3); and active chromatin marks (H3K27ac, H3K4me3, H3K9ac) are maintained across all analyzed TB lineages **(Fig 2b)**. These observations indicate *SUPYN* expression persists through the TB differentiation process. We next mined single-cell RNA-seq data generated from placenta at multiple developmental stages. After classifying cell clusters based on known markers, we found *SUPYN* and *ASCT2* expression was relatively high in cytotrophoblasts (CTB) and extra-villous trophoblasts (EVTB), but also detectable in syncytiotrophoblasts (STB) **(Fig 2c-f, fig S6)**. To confirm these transcriptomic observations, we performed immunostaining of SUPYN in preimplantation embryos as well as in second and third trimester placentas. In early embryos, SUPYN is primarily detectable in the trophectoderm, which will give rise to the placenta, and in some OCT4-expressing cells, which mark pluripotent stem cells of blastocysts **(Fig 2g, fig S7)**. In the placenta, SUPYN is widely expressed in STB and potentially CTB of second trimester placental villi **(Fig 2h, fig S8)**. The combined transcriptome, genome regulatory data and immunohistochemical staining firmly establish that *SUPYN* is expressed throughout human fetal development (**Fig 2a, 2e, fig S6-8)**. These analyses also indicate that *SUPYN* transcription consistently initiates from its ancestral LTR promoter **(Fig 2b, fig S5b, S13)**.

**Figure 2:**
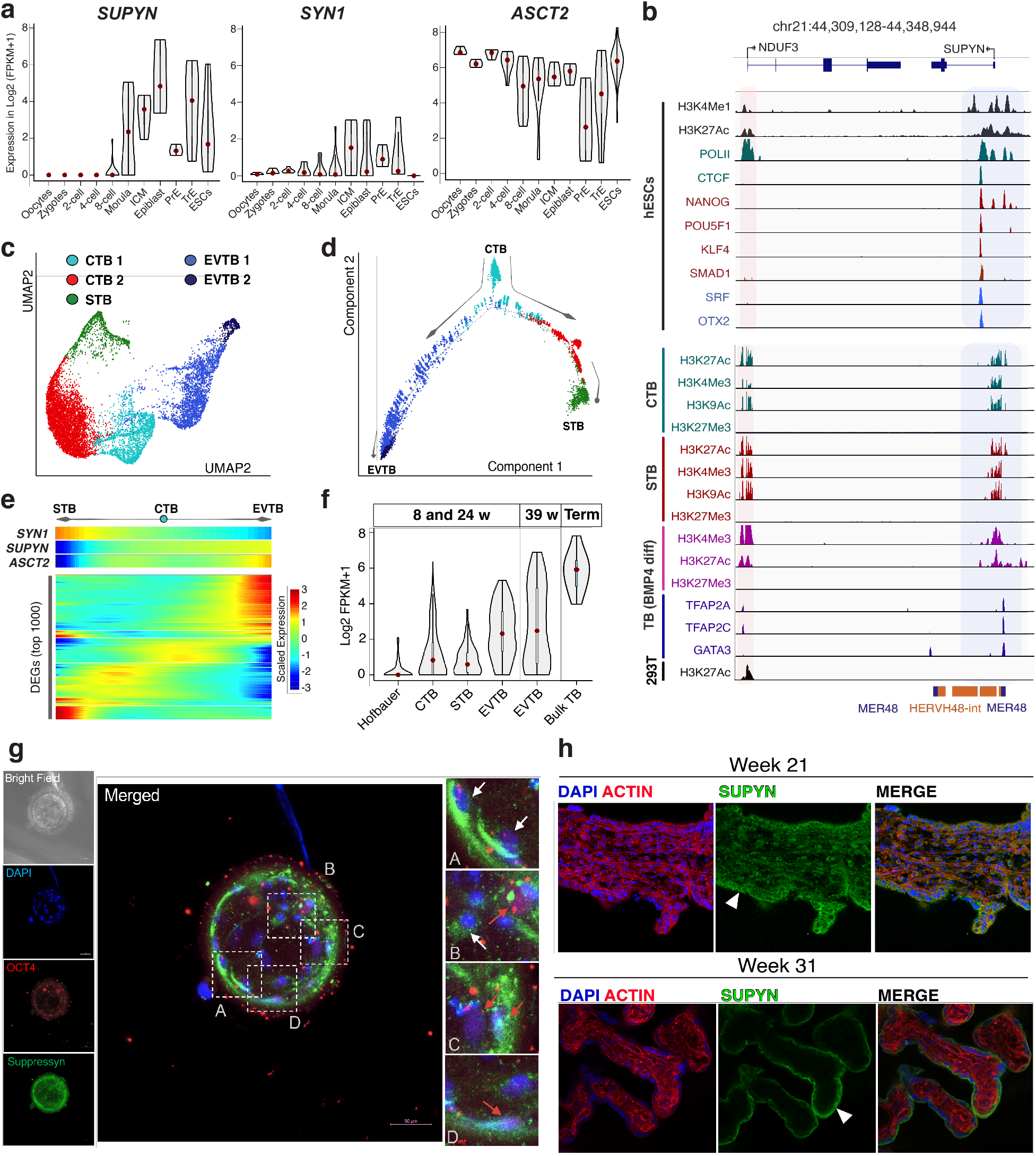
Pluripotency and placentation regulatory factors drive SUPYN expression during fetal development. **(a)** Violin plots summarizing *SUPYN, SYN1* and *ASCT2* expression in human preimplantation embryo and ESC single-cell RNA-seq data. **(b)** Genome browser view of the regulatory elements surrounding the *SUPYN* locus in hESCs, cyto- (CTB), syncytio- (STB), BMP4-differentiated trophoblasts (TB), and 293T cells. ChIP-seq profiles are shown for indicated transcription factors and histone modifications with shaded area highlighting regions of active chromatin. **(c)** UMAP visualization of TB cell clusters, shown in figure S6. **(d)** Monocle2 pseudotime analysis of cell clusters in (c) illustrates the developmental trajectory of CTBs that give rise to STB and EVTB respectively. Color codes in (c) and (d) denote cell identity. **(e)** Heatmap represents the top 1000 differentially expressed genes (row) of single cells (column) and are sorted according to pseudotime analyses in (c) and (d). Cells are ordered according to the pseudotime progression of CTB (middle) to STB (left) and EVTB (right). SYN1, SUPYN, and ASCT2 were fetched from the heatmap below. **(f)** Violin plots denote single-cell *SUPYN* expression in multiple placental-cell lineages throughout indicated pregnancy stages. **(g**,**h)** Confocal microscopy of human preimplantation embryos (g) and placental villus explants (h) stained for SUPYN (green), OCT4 (red), ACTIN (red) and DAPI (blue). In (g), Trophectorderm (white arrows) and OCT4-expressing (red arrows) cells are highlighted in subpanels (A-C). In (h), STBs are marked by arrowheads.

The expression profile of SUPYN is consistent with the hypothesis that it could protect the developing embryo and germline against RDR infection. To test this hypothesis, we first examined whether human placenta-derived Jar and JEG3 cell lines, and H1 hESC, which robustly express SUPYN and ASCT2 **(Fig 1a, 2a; fig S9, S10)** (*10, 13*) are resistant to RDR env-mediated infection. We generated HIV-EGFP viral particles pseudotyped with either the feline RD114 env (HIV-RD114) or the glycoprotein G of vesicular stomatitis virus (HIV-VSVg), which uses the LDL receptor (*14*), to monitor the level of infection in cell culture based on EGFP expression **(Fig 3a, fig S11)**. These experiments revealed that Jar, JEG3, and H1 cells were susceptible to HIV-VSVg, as previously reported (*15, 16*), but highly resistant to HIV-RD114 infection **(Fig 3b, c)**. By contrast, 293T cells, which do not express *SUPYN* **(fig S5a)**, were similarly susceptible to infection by HIV-RD114 and HIV-VSVg **(Fig 3b, c)**. To test whether SUPYN contributes to the HIV-RD114 resistance phenotype, we repeated these infection experiments in Jar cells engineered to stably express short hairpin RNAs (shRNA) depleting ∼80% of *SUPYN* mRNA **(fig S10a)**. Depletion of *SUPYN* in Jar cells resulted in a significant increase in susceptibility to HIV-RD114 infection but not HIV-VSVg **(Fig 3d**). To confirm that SUPYN expression confers RD114 resistance and control for potential off-target effects of *SUPYN*-targeting shRNAs, we transfected Jar-shSUPYN cells with two shRNA-resistant, HA-tagged SUPYN rescue constructs and examined their susceptibility to HIV-RD114 infection. Briefly, the shRNA-targeted SUPYN signal peptide sequence was replaced with a luciferase (SUPYN-lucSP) or modified signal peptide sequence (SUPYN-rescSP) that disrupts shRNA-binding but retains codonidentity (see Methods). Transfection with either SUPYN-rescSP or SUPYN-lucSP restored a significant level of resistance to HIV-RD114 infection **(Fig 3e)**. Western blot analysis of transfected cell lysates showed SUPYN-rescSP was more abundantly expressed than SUPYN-lucSP **(Fig 3f)**, which may account for the stronger HIV-RD114 resistance conferred by the former **(Fig 3e)**. These results suggest that SUPYN expression is at least in part responsible for preventing Jar cells from RD114env-mediated infection.

**Figure 3:**
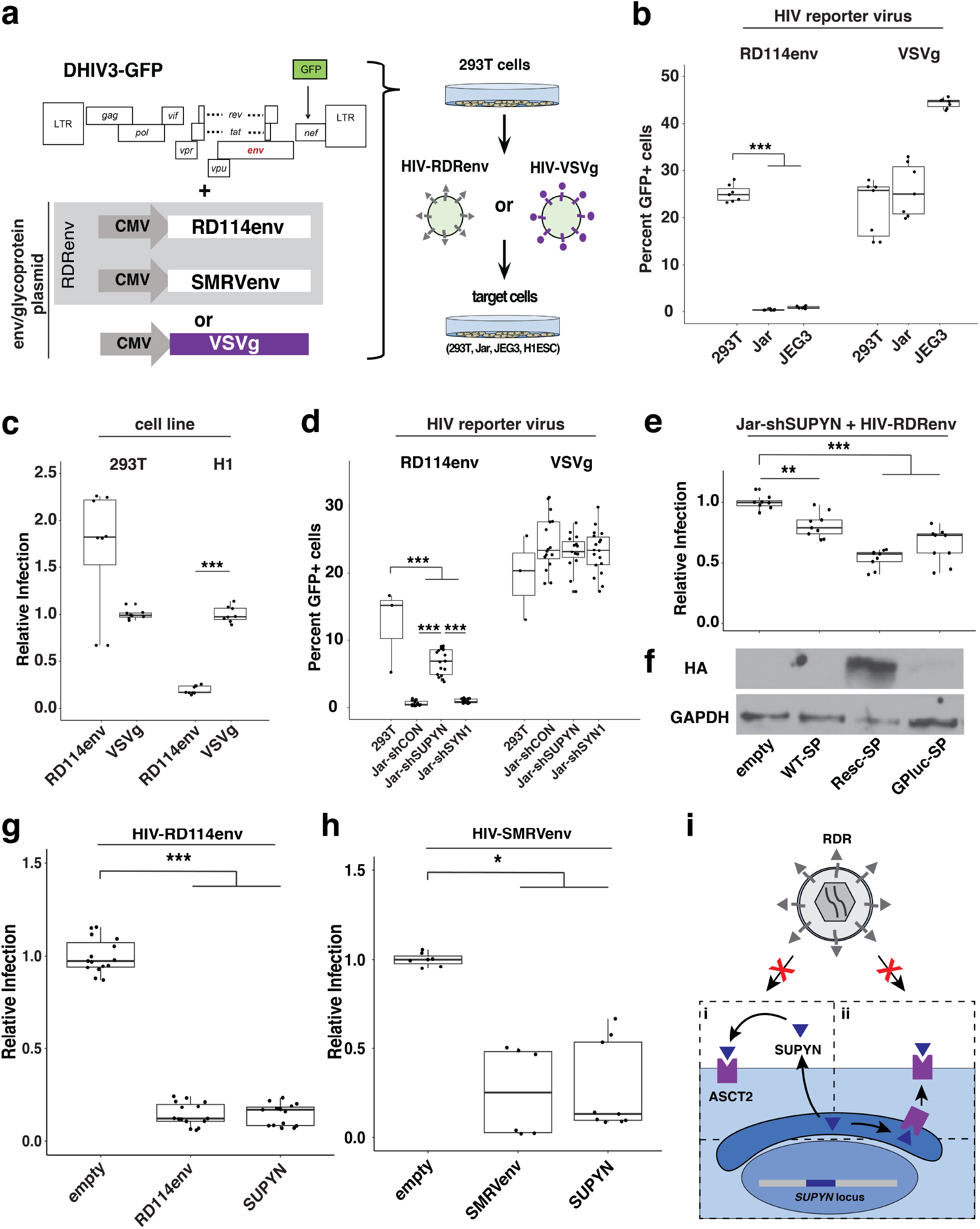
SUPYN confers resistance to RD114 env-mediated infection. **(a)** HIV-glycoprotein reporter virus production and infection assay approach (see Methods). **(b)** Proportion of GFP+ 293T, JEG3, and Jar cells infected with indicated reporter viruses. **(c)** Relative infection rate of 293T and H1-ESCs was normalized to mean proportion of HIV-VSVg-infected cells. **(d)** Proportion of GFP+ 293T and shSUPYN-Jar cells infected with indicated reporter virus. **(e)** Relative infection rates of shSUPYN-Jar cells transfected with HA-tagged wild-type (WT-SP), rescue signal peptide (Resc-SP), or luciferase signal peptide (GPluc-SP) encoding SUPYN overexpression constructs. (n 3 with 2 technical replicates; ***adj. *p < 0.001*; **adj. *p < 0.01*; ANOVA with Tukey HSD). **(f)** Western Blot analysis (HA, GAPDH) of Jar-shSUPYN cell lysates transfected with indicated SUPYN overexpression constructs. **(g**,**h)** Relative infection rates of 293T cells transfected with unmodified SUPYN (g, h), RD114env (g), or SMRVenv overexpression plasmids (h) and subsequently infected with indicated reporter viruses. **(i)** Model of SUPYN-dependent RDR infection restriction. SUPYN likely binds ASCT2 following secretion (i) or within the secretory compartment (ii).

To test if SUPYN expression alone is sufficient to confer protection against RD114env-mediated infection, we transfected 293T cells with unmodified or HA-tagged SUPYN overexpression constructs and subsequently infected with HIV-RD114 and HIV-VSVg respectively. As a positive control, we also transfected 293T cells with an unmodified RD114env overexpression construct, which is predicted to confer resistance to HIV-RD114. Expression of either SUPYN or RD114env resulted in ∼80% reduction in the level of HIV-RD114 infection **(Fig 3g; fig S12a)** but had no significant effect on HIV-VSVg infectivity **(fig S12b)**. Taken together, our knockdown and overexpression experiments indicate SUPYN expression is sufficient to confer resistance to RD114env-mediated infection.

Our RD114env-specific resistance phenotype **(Fig 3g, fig S12a, b)** suggests SUPYN restricts viral entry via a mechanism of receptor interference. If so, this protective effect should extend to infection mediated by other RDRenv (*11, 17, 18*). To test this, we generated HIV-GFP reporter virions pseudotyped with Squirrel Monkey Retrovirus env (HIV-SMRVenv) (*11*) **(Fig 3h)** and infected 293T cells previously transfected with SUPYN, SMRVenv (as a positive control) or an empty vector. Cells expressing SUPYN or SMRVenv showed an ∼80% reduction of HIV-SMRVenv infected cells **(Fig 3h)**. Thus, SUPYN can restrict infection mediated by multiple RDRenv.

Another prediction of RDR restriction via receptor interference is that it should be a property of env proteins recognizing ASCT2, such as SUPYN, but not those binding other cellular receptors. Consistent with this prediction and previous work with overexpressed SYN1 (*19*), expressing HA-tagged SUPYN and SYN1 strongly restricted HIV-RD114, while HA-tagged env from amphotrophic murine leukemia virus (aMLV) or human endogenous retrovirus H (HERVH), neither of which are expected to interact with ASCT2 (*20*–*22*), had no effect on HIV-RD114 nor HIV-VSVg infection in 293T cells **(fig S12a, b)**. Importantly, all tested env proteins were expressed at comparable levels **(fig S12c)**. Furthermore, we observed that SUPYN overexpression did not significantly alter the level of ASCT2 protein expression in 293T cells **(Fig S12c)**. This result suggests that if SUPYN acts by receptor interference, its interaction with ASCT2 does not result in receptor degradation, which is consistent with some instances of receptor interference (*23*–*25*). In agreement with previous observations (*5*), we noted that *SUPYN* knockdown in Jar cells resulted in the selective loss of a putative non-glycosylated isoform of ASCT2 **(fig S10b)**. We speculate that the presence of SUPYN-dependent non-glycosylated ASCT2 may be the result of SUPYN sterically interfering with the glycosylation machinery within the secretory pathway. Collectively, these observations converge on the model that SUPYN restricts against RDR infection through receptor interference by two possible mechanisms that are not mutually exclusive: SUPYN may bind ASCT2 within the secretory pathway or extracellularly following secretion **(Fig 3i)**.

To gain insights into the evolutionary emergence of *SUPYN* antiviral activity, we used comparative genomics to investigate the origin and functional constraint acting on *SUPYN*. We found that the HERVH48 provirus from which *SUPYN* is derived is present at orthologous position across the genomes of all available hominoids and most Old World monkeys (OWM), but is absent in other primate lineages **(Fig 4a, fig S13)**. Thus, the provirus that gave rise to *SUPYN* inserted in the common ancestor of catarrhine primates ∼27-38 million years ago (*26*) **(Fig 4a)**. We next examined whether *SUPYN* ORFs have evolved under functional constraint during primate evolution. The *SUPYN* ORF is almost perfectly conserved in length across hominoids, but not in OWM where some species display frameshifting and truncating mutations **(Fig 4a, fig S14, S15)**, suggesting *SUPYN* evolved under different evolutionary regimes in hominoids and OWMs. To test this idea, we analyzed the ratio (*ω*) of nonsynonymous (dN) to synonymous (dS) substitution rates using codeml (*27*), which provides a measure of selective constraint acting on codons. Log-likelihood ratio tests comparing models of neutral evolution with selection indicate *SUPYN* evolved under purifying selection in hominoids (*ω* = 0.38; *p* = 1.47E-02) but did not depart from neutral evolution in OWMs (*ω* = 1.44; *p* = 0.29) **(Fig 4a)**. These results indicate that *SUPYN* antiviral activity may be conserved across hominoids but not in OWM. To assess this, we transfected 293T cells with HA-tagged overexpression constructs for the orthologous *SUPYN* sequences of two hominoid species (chimpanzee, siamang), and five OWM species (African green monkey, pigtailed macaque, crab-eating macaque, rhesus macaque, olive baboon) and challenged these cells with HIV-RD114. Both chimpanzee and siamang SUPYN proteins displayed antiviral activity with potency comparable to or greater than human SUPYN, respectively **(Fig 4b)**. By contrast, of the five OWM, only African green monkey SUPYN exhibited a modest but significant level of antiviral activity **(Fig 4b, c)**. The lack of restriction activity for some of the OWM proteins may be attributed to their relatively low expression level in these human cells **(Fig 4b)** and/or their inability to bind the human ASCT2 receptor due to *SUPYN* sequence divergence **(fig S14, S15)**. To further trace the evolutionary origins of SUPYN antiviral activity, we expressed ancestral *SUPYN* sequences phylogenetically reconstructed for the hominoid and OWM ancestors **(Fig S15)** and assayed their antiviral activity in 293T cells. Both ancestral proteins were expressed at levels comparable to human and green monkey SUPYN and exhibited strong restriction activity **(Fig 4c)**. These data indicate that SUPYN antiviral activity against RDRenv-mediated infection is an ancestral trait preserved over ∼20 million years of hominoid evolution, but apparently lost in several OWM lineages.

**Figure 4:**
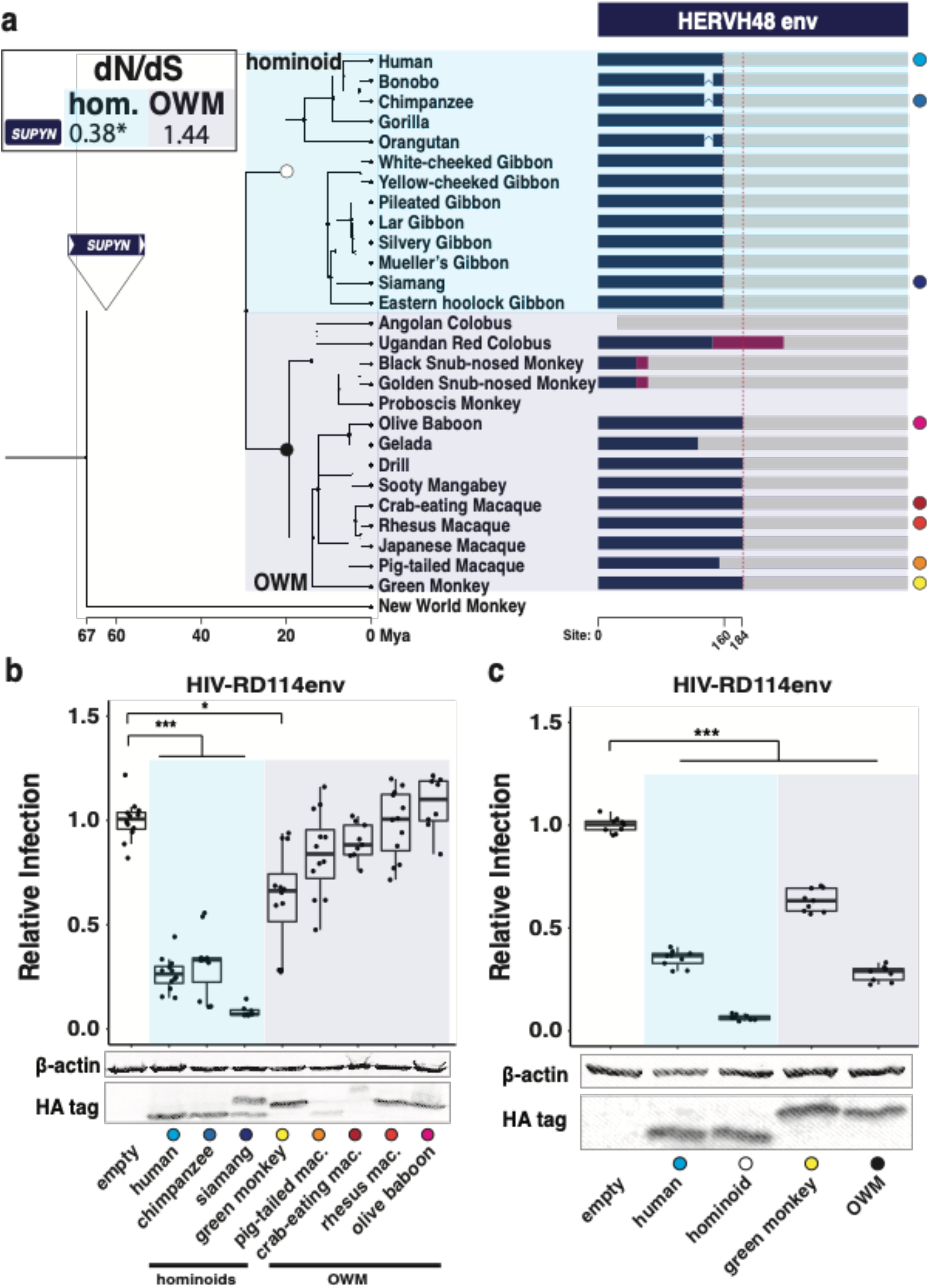
SUPYN emerged in a Catarrhine ancestor and has conserved antiviral activity in Hominoids. **(a)** Consensus primate phylogeny with cartoon representation of *SUPYN* ORFs (blue box). Magenta boxes represent frameshifts in *SUPYN* ORFs. Red dashed lines denote conserved premature stop codon positions. Grey bars represent degraded HERVH48env sequence. Labeled triangles denote ancestral lineage where HERVH48env was acquired. Colored circles indicate species used in **b** and **c**. *SUPYN* dN/dS value is shown in box (\**p < 0.05*; LRT). **(b, c)** Relative infection rates and Western Blot of 293T cells transfected with primate (**b**) or ancestral (**c**) SUPYN-HA constructs and subsequently infected with HIV-RD114env are shown. Relative infection rates were determined by normalizing GFP+ counts to empty vector. (n 3 with 2 technical replicates ***adj. *p < 0.001*; *adj. *p < 0.05*; ANOVA with Tukey HSD)

In this study, we show that the human genome harbors a large pool of novel and previously annotated envORF sequences (*6, 7*) that are dynamically expressed, particularly in the germline, early embryonic development, and in response to immune stimulation and infection. Because envORFs are predicted to encode proteins with the ability to bind receptors used by their ancestral retroviruses and possibly many contemporary exogenous viruses, we propose that they constitute a reservoir of potential antiviral factors. As a proof of principle, we established that one of these proteins, *SUPYN*, is both necessary and sufficient to confer resistance to RDRenv-pseudotyped reporter virus infection in human cell culture. We further determined that SUPYN expression in the human preimplantation embryo and throughout placental development is driven by a tightly regulated promoter derived from the LTR of its ancestral provirus. These results suggest that the complex cis-regulatory activities encoded by retroviral LTRs facilitate ERV gene cooption (*1, 2*) The prominent expression of envORF, including SUPYN, in the developing germ line suggests that they may have repeatedly acted as a barrier against the vertical propagation of the cognate endogenous retroviruses by reinfection (*28*). This may explain why most HERV families appear to have propagated in the genome primarily via retrotransposition rather than reinfection (*29*). Under this model, we predict the adaptive benefits of envORFs to be generally transient, unless they have broader antiviral activity or gain additional cellular function. This conjecture may explain why *SUPYN* has been preserved by natural selection throughout hominoid evolution; not only may *SUPYN* have shielded the early embryo and nascent germline from the persistent threat of RDRs but also against the adverse effects of SYN1-mediated infections and syncytalization of the developing placenta (*5, 10, 30*). In conclusion, this study provides evidence that truncated envelope-derived peptides encoded in the human genome can evolve and maintain antiviral activity. It is possible that our genome holds many other retrovirus-derived proteins with protective effects against viral infection.

## Methods

### RNA-seq analyses

We mined published single cell transcriptome datasets (scRNA-seq) of human pre-implantation embryos isolated at developmental stages ranging from oocyte to blastocyst (*31*), primordial germ cells (*32*), human placenta (*33*–*35*), neuronal differentiation (*36*), haematopoietic stem cells (*37*), pancreas (*38*), prefrontal cortex (*39*). Prior to analyzing the transcriptomic datasets, we first recompiled the reference genome annotations using known gene (genecode V19) (*40*), and *envORF* sequences to generate a reference transcriptome in fasta format. To guide the transcriptome assembly, we converted this updated genome annotation to gtf format, which was subsequently utilized for our expression quantifications. Reads were mapped to the curated human genome (hg19) with STAR (*41*) using the following settings --*alignIntronMin 20 -- alignIntronMax 1000000 --chimSegmentMin 15 --chimJunctionOverhangMin 15 -- outFilterMultimapNmax 20*. Only uniquely mapped reads were considered for expression calculations. Gene level counts were obtained using *featureCounts* (*42*) run with RefSeq annotations. Gene expression levels were calculated at <u>T</u>ranscript <u>P</u>er <u>M</u>illion (TPM) from counts mapped over the entire gene (defined as any transcript located between <u>T</u>ranscription <u>S</u>tart <u>S</u>ite (TSS) and <u>T</u>ranscription <u>E</u>nd <u>S</u>ite (TES)). Only genes and cells that met the following criteria were included in this analysis: (1) each cell must express at least 5,000 genes; (2) each gene must be expressed in at least 1% of cells; (3) each gene must be expressed with log2 TPM >1. We clustered cells meeting these criteria using the default parameters of the Seurat (*43*) package (v3.1.1) implemented in R (v3.6.0). Seurat applies the most variable genes to get top principal components that are used to discriminate cell clusters in tSNE or UMAP plots. We set Seurat to use 10 principal components in this cluster analysis. For the placental scRNAseq data **(Fig 2, Fig S5)**, the 2000 most differentially expressed genes were used to define cell clusters. Major clusters corresponding to CTB, STB, EVTB, macrophages, and stromal cells were identified based on the expression of known marker genes. Monocle2 (*44*) was used to perform single-cell trajectory analysis and cell ordering along an artificial temporal continuum. The top 500 differentially expressed genes were used to distinguish between CTB, STB and EVTB cell populations. The transcriptome from each single cell represents a pseudo-time point along an artificial time vector that denotes the progression of CTB to STB or EVTB respectively.

Data generated on the 10X Genomics scRNA-seq platforms were processed in the following way. Normalized counts and cell-type annotations were used as provided by the original publications. Seurat was used for filtering, normalization, and cell-type identification. The following data processing steps were performed: (1) Cells were filtered based on the criteria that individual cells must have between 1,000 and 5,000 expressed genes with a count ≥1; (2) cells with more than 5% of counts mapping to mitochondrial genes were filtered out; (3) data was normalized by dividing uniquely mapping read counts (defined by Seurat as unique molecular identified (UMI)) for each gene by the total number of counts in each cell and multiplying by 10,000. These normalized values were then natural-log transformed. Cell types were defined by using the top 2000 variable features expressed across all samples. Clustering was performed using the “FindClusters” function with largely default parameters; except resolution was set to 0.1 and the first 20 PCA dimensions were used in the construction of the shared-nearest neighbor (SNN) graph and the generation of 2-dimensional embeddings for data visualization using UMAP. Cell types were assigned based on the annotations provided by the original publication.

Bulk RNAseq datasets generated from the GTEx consortium (*45*), placenta (*46*), 293T (*47, 48*), and immune (*49, 50*) cells were processed as described above. Briefly, reads were mapped with STAR and uniquely mapped reads were counted with featureCounts. We restricted the visualization to only *envORFs* that were expressed (Log2 TPM > 1) in at least one of the analyzed or compared samples (see the codes submitted on GitHub).

### ChIP-seq, DNAse-seq and ATAC-seq data analysis

Various ChIP-seq datasets representing histone modifications and transcription factors in human embryonic stem cells and their differentiation were retrieved from (*51, 52*). We obtained the H3K27Ac for CTB to STB primary culture (*53*), H3K4Me1 for trophoblasts (*54*), H3K4Me3, H3K27Me3 for differentiated trophoblasts (*55*), and GATA2/3, TFAP2A/C (*55*) ChIP-seq datasets in raw fastq format. DNAse-seq and ATAC-seq datasets were retrieved from references (*56*) and (*57*) respectively.

Reads from the above-described datasets were aligned to the hg19 human reference genome using Bowtie2 (*58*) run in *--very-sensitive-local* mode. All reads with MAPQ < 10 and PCR duplicates were removed using Picard and *samtools* (*59*). Peaks were called by MACS2 (*60*) (https://github.com/macs3-project/MACS) with the parameters in narrow mode for TFs and broad mode for histone modifications keeping FDR < 1%. ENCODE-defined blacklisted regions (*61*) were excluded from called peaks. We then intersected these peak sets with repeat elements from hg19 repeat-masked coordinates using bedtools *intersectBed* (*62*) with a 50% overlap. To visualize over Refseq genes (hg19) using IGV (*63*), the raw signals of ChIP-seq were obtained from MACS2, using the parameters: *-g hs -q 0*.*01 -B*. The conservation track was visualized through UCSC genome browser (*13*) under net/chain alignment of given non-human primates (NHPs) and merged beneath the IGV tracks.

### Cell culture

293T cells were cultured in DMEM (GIBCO, 11995065) containing 10% Fetal Bovine Serum (FBS) (GIBCO, 10438026). Jar cells (provided by Carolyn Coyne) were cultured in RPMI (GIBCO, 11875093) containing 10% FBS. JEG3 cells were cultured in MEM (GIBCO, 11095080) containing 10% FBS. Culture medium for these cell lines was supplemented with sodium pyruvate (GIBCO, 11360070), glutamax (GIBCO, 35050061), and Penicillin Streptomycin (GIBCO, 15140122) according to manufacturer specifications. H1-ESCs (*64*) (obtained from WiCell) were grown on Matrigel (Corning, 356277) coated plates in MTESR+ (Stemcell, 05825) growth-media and sub-cultured using Accutase (Innovative Cell Technologies, AT-104) and MTESR+ supplemented with CloneR (Stemcell, 05888). All cell lines were cultured at 37°C and 5% CO2.

### Human embryos

Prior to starting the project (documentation available upon request), all the procedures to obtain human pre-implantation embryos were approved by local regulatory authorities and the Spanish National Embryo steering committee. After the signing of informed consent documents by parents, all human pre-implantation embryos were donated and fully anonymized. The embryos used in this study were cryopreserved sibling embryos after successful In Vitro Fertilization pregnancies.

### Culturing of human pre-implantation embryos

Human pre-implantation embryos at several stages of development (from +2 to +6 days post-fertilization) were thawed and allowed to grow. Only embryos reaching the blastocyst stage and showing viability in most individual cells were selected for further analyses. Cryopreserved human embryos were thawed using thawing media from Vitrolife (G-1 v.5 and G-2 v.5 media), and were classified based on morphological criteria using inverted Hoffman optical microscopy. More than 50% of thawed embryos showed signs of cell division.

### Vector cloning

DHIV3-GFP, phCMV-RD114env, psi(-)-amphoMLV plasmids were provided by Vicente Planelles (University of Utah). pCGCG-SMRVenv plasmid was provided by Welkin Johnson (Boston University). psPAX2 and pVSVg plasmids were provided by John Lis (Cornell University). The following cloning approaches were performed using primers and constructs described in Supplementary Table 3. HERVH1env ORF was PCR-amplified using Q5 polymerase (NEB, M0491L) from HeLa and 293T genomic DNA respectively and cloned into a TOPO vector (ThermoFisher, 450245).

To generate stable *SYN1* and *SUPYN* knock-down cell lines, pHIV lentiviral constructs containing shRNAs targeting *SYN1* and *SUPYN* respectively were cloned using the following strategy. The shRNA encoded in pHIV7-U6-shW3, generously provided by Lars Aagaard (Aarhus University), targets *SYN1* (*65*). *SUPYN*-targeting shRNAs were designed using siRNA sequences employed by Jun Sugimoto^4^ as a template. pHIV7 lentiviral constructs were cloned using the pHIV7-U6-shW3 plasmid (*65*) as a template. pHIV7-U6-shSup-cer, pHIV7-U6-shSup-puro, pHIV7-U6-shC-cer, pHIV7-U6-shC-puro, pHIV7-U6-shSyn1-cer, pHIV7-U6-shSyn1-puro were generated using a Gibson assembly approach. To replace the native GFP marker of pHIV7-U6-shW3 with a Cerulean reporter or puromycin resistance marker, we digested pHIV7-U6-shW3 with NheI (NEB, R3131S) and KpnI (NEB, R3142S). This digest resulted in the production of three DNA fragments: pHIV7 backbone, GFP-, and WPRE-containing fragments. We separately PCR amplified each selection marker and WPRE containing pHIV7 fragment. InFusion cloning was then used to ligate the digested pHIV7 backbone to the Cerulean or puromycin cassette and WPRE containing PCR product. shRNAs were cloned into the pHIV7-Cerulean/puromycin transfer construct previously digested with NotI (NEB, R0189S) and NheI. U6-promoter containing shRNA cassettes and the CMV promoter driving marker cassette expression were PCR amplified and subsequently InFusion cloned into the NotI/NheI digested pHIV7-cerulean/puromycin backbone.

All pHCMVenv and SUPYN expression constructs, described in this study, were generated as follows: HA-tagged and untagged ORFs with pHCMV homologous overhanging sequence were either PCR amplified using Q5 polymerase (NEB, M0491S) or synthesized (IDT) (see **Table S2**), and cloned into EcoRI (NEB, R3101T) digested pHCMV backbones using the InFusion cloning kit (Takara Bio, 638920). To generate siRNA-resistant SUPYN rescue constructs, we replaced the native signal peptide sequence^4^ (which is targeted by siRNAs used in this study) with (1) a *Gaussia princeps* luciferase SP (SUPYN-lucSP) (*66, 67*) and (2) a shSUPYN resistant SUPYN rescue construct (SUPYN-rescSP) in which the codons were modified to retain the codon identity but disrupt siRNA binding.

### Antibodies

All antibodies used in this study are commercially available. α-GAPDH (D4C6R, D16H11), α-βactin (D6A8), α-HA (C29F4), α-ASCT2 (V501) primary antibodies were purchased from Cell Signaling Technology. α-SUPYN and αOCT4 primary antibodies were purchased from Phoenix Pharma (H-059-052) and Santa Cruz Biotechnology respectively. α-Mouse (#7076) and α-Rabbit (#7074) HRP conjugated secondary antibodies were purchased from Cell Signaling Technology. IRDye secondary antibodies were purchased from Licor (925-32211, 925-68072, 925-32210, 925- 68073). Alexa-fluor conjugated secondary antibody was purchased from Invitrogen.

### Western Blot

Whole cell extracts from cultured cell lines were prepared using 1x GLO lysis buffer (Promega, E266A). One third volume of 4x Laemli buffer was added to one volume whole cell extract samples, then incubated at 95°C for 5 minutes, and sonicated for 15 minutes at 4°C (amplitude 100; pulse interval 15 seconds on, 15 seconds off). Approximately 30 ug of protein were separated by SDS-PAGE (BioRad, 1610175), transferred to PVDF membrane (BioRad, 1620177), blocked according to antibody manufacturers specification, and incubated overnight in appropriate primary antibody then incubated in IRDye or peroxidase conjugated goat anti-mouse or anti-rabbit secondary antibodies for 1 hour at room temperature. Protein was then detected using ECL reagent (BioRad, 1705061) or the Licor Odyssey imaging system.

### IF microscopy

#### For placenta

Human second trimester placental tissue that resulted from elective terminations was obtained from the University of Pittsburgh Health Sciences Tissue Bank through an honest broker system after approval from the University of Pittsburgh Institutional Review Board (IRB) and in accordance with the University of Pittsburgh’s tissue procurement guidelines. Tissue was excluded in cases of fetal anomalies or aneuploidy. Third trimester placental tissue was obtained through the Magee Obstetric Maternal & Infant (MOMI) Database and Biobank after approval from the University of Pittsburgh IRB. Women who had previously consented for tissue donation and underwent cesarean delivery were included. Placental tissues were fixed in 4% PFA (in 1x PBS) for 30 minutes, permeabilized with 0.25% Triton X-100 for 30 minutes (on a rocker), washed with 1x PBS and then incubated with primary anti-Suppressyn antibody at 1:200 in 1x PBS for 2-4 hours at room temperature. These samples were incubated with Alexa-fluor conjugated secondary antibody (Invitrogen) diluted 1:1000 and counterstained with actin. DAPI was included in our PBS and then mounted in Vectashield mounting medium with DAPI (Vector Laboratories, H-1200).

#### For embryo

Pre-implantation embryos at the blastocyst stage were washed with 1x PBS (Invitrogen) and fixed for 15 minutes using fresh 4% (w/v) paraformaldehyde (Sigma) at room temperature. After fixation, embryos were washed twice on a petri dish using 1x PBS containing 0.1% (v/v) Triton X-100 (Sigma). Next, embryos were permeabilized on a lo-bind 2ml Eppendorf tube using 1x PBS containing 0.5% (v/v) Triton X-100 for 24 hours at 4°C. After permeabilization, embryos were incubated for 24 hours at 4°C in blocking solution [1x PBS containing 0.1% (v/v) Triton X-100, 1% (v/v) Tissue Culture Grade DMSO (Sigma), and 5% (v/v) Normal Goat Serum (Abcam)]. Next, embryos were incubated with the primary antibodies [1:200 anti-Rabbit Suppressyn (Phoenix Pharmaceuticals, Inc), and 1:200 anti-Goat OCT4 (Santa Cruz Biotechnology, Inc)] in fresh blocking solution for 24 hours at 4°C. After 30 minutes at room temperature, embryos were washed twice for 5 minutes using 1x PBS containing 0.1% (v/v) Triton X-100, and subsequently incubated with secondary antibodies in blocking solution containing DAPI (Thermo Scientific) [1:1000 anti-Rabbit Alexa 488 (Life Technologies), and 1:1000 anti-Goat Alexa 555 (Life Technologies)] for 24 hours at 4°C in the dark. Next, embryos were washed twice for 5 minutes using 1x PBS containing 0.1% (v/v) Triton X-100 and mounted in a μ-Slide 8 well (Ibidi®) using 1xPBS and avoiding the generation of air bubbles.

A confocal Zeiss LSM 710 device (Genyo, Spain) was used to process the slides, using a 63X objective, zoom 0.6x, with a 1024×1024 pixels resolution (corresponding to 224.70 µm x 224.70 µm). Fluorescence channels were recorded independently to avoid artifacts, and microscope images shown are maximum projections of several images (2-3 or more). As negative controls, embryos were also incubated with only the secondary antibody.

### Virus production

Low passage 293T cells were used to produce lentiviral particles. DHIV3-GFP and env-expression plasmids were co-transfected at a mass ratio of 2:1 using lipofectamine 2000 (ThermoFisher, 11668030). shRNA encoding lentiviral particles were produced by co-transfecting pHIV7, psPAX2, pVSVg according to BROAD institute lentiviral production protocol (https://portals.broadinstitute.org/gpp/public/resources/protocols) using Lipofectamine 2000. Growth media was replaced on transfected cells after overnight incubation. At 72 hours post-transfection, virus containing supernatant was harvested, centrifuged to remove cell debris, filtered through a 0.45 um pore filter, and stored at −80°C.

### Infection Assays

293T cells were transfected with env-overexpression constructs using Lipofectamine 2000 and incubated 24 hours. Transfected cells were infected with reporter virus by applying virus (HIV-RD114env, HIV-VSVg, HIV-SMRVenv) stocks in the presence of polybrene (Santa Cruz Bio, sc-134220) at a final concentration of 4 ug/mL. After 6-8 hours, virus stock was replaced with fresh growth media. Infected cells were maintained for 72 hours, replacing media when necessary, and harvested with trypsin. Detached cells were suspended in fresh growth media, strained and analyzed by flow cytometry. For the H1-ESC infection experiment, relative infection rates were calculated by normalizing the percent GFP+, HIV-RD114env infected cells to the percent GFP+ HIV-VSVg infected cells. For env/SUPYN overexpression experiments, relative infection rates were calculated by normalizing the percent GFP+ env/SUPYN-transfected cells to the percent GFP+ empty vector transfected cells. ANOVA with Tukey HSD tests were implemented in R (v3.6.3).

### Placental cell shRNA transduction

Placenta-derived cell lines were treated with pHIV-shRNA-virus-containing supernatant and incubated for 72 hours as described in Infection Assays. Cerulean positive cells were sorted using the BD FACS Aria cytometer. Cells transduced with puroR cassette were treated with Puromycin (GIBCO, A1113802) at a final concentration of 3.5 ug/mL for 7 days, then cultured in regular growth media.

### RT-qPCR

RNA was isolated from cultured cells using the RNeasy Mini Kit (Qiagen, 74104) and an on column dsDNAse digestion (Qiagen, 79254) was performed. 1-3 ug of total RNA were used to generate cDNA with the maxima cDNA synthesis with dsDNAse kit (ThermoFisher, K1681). qPCR reactions were performed using the LC480 Instrument with Sybr Green PCR master mix (Roche, 04707516001) according to manufacturer’s protocol and using primers indicated in Supplementary Table 2. Gene expression was then quantified using the ΔΔCT method (*68*). 18S expression was used as a reference housekeeping gene. Wilcox rank sum tests were performed using R (v3.6.3).

### Envelope evolutionary sequence analyses

Orthologous *SUPYN, SYN1*, and *SYN2* sequences were extracted from the 30-species MULTIZ alignment (*13*) and formatted for sequence alignment using the phast package (*69*). These and additional syntenic *SUPYN* and *SYN2* open reading frame sequences were validated/identified by BLASTn (*70*) search with default settings of publicly available Catarrhine primate genomes (ncbi.nih.gov). Mariam Okhovat of the Carbone Lab (Oregon Health and Science University) generously provided BAM files containing read alignment information for *SUPYN, SYN1*, and *SYN2* generated from whole genome sequencing of *Hoolock leuconedys* (Hoolock Gibbon), *Symphalangus syndactylus* (Siamang), *Hylobates muelleri* (Müller’s Gibbon), *Hylobates lar* (Lar Gibbon), *Hylobates moloch* (Silvery Gibbon), *Hylobates pileatus* (Pileated Gibbon), and *Nomascus gabriellae* (Yellow-cheeked Gibbon) (*71*). Where multiple individuals were sequenced, a consensus sequence was generated using samtools (*59*) and JalView (*72*).

To perform *dN/dS* analyses, orthologous env sequences (>90bp length) encoding the mature sequence downstream of the signal peptide cleavage site, were aligned using MEGA7 (*73*) and manually converted to PHYLIP format. A Newick tree was generated based on this alignment using the maximum likelihood algorithm implemented in MEGA7. *Codeml*, implemented in the PAML package, was run to calculate *dN/dS* values and log likelihood (LnL) scores generated under models M0, M1, M2, M7 and M8 (*27*). Chi-square tests comparing LnL scores generated under models of neutral evolution and selection were performed.

We used two approaches to reconstruct ancestral hominoid and OWM *SUPYN* sequences. First, we reconstructed ancestral *SUPYN* sequences using the majority rule consensus sequence (calculated in JalView) of the hominoid and OWM clade respectively. At positions where nucleotide identity was ambiguous, the dominant nucleotide identity in the neighboring clade was used as a tiebreaker. These sequences were used for our infection assays shown in **Fig 4h**. We also employed a maximum likelihood approach using the *baseml* program, implemented in PAML (*27*). We reconstructed ancestral *SUPYN* sequences using the hominoid species, shown in **Fig 1**, and the 6 OWM monkeys with the most complete *SUPYN*-coding open reading frame (olive baboon, drill, crab-eating macaque, rhesus macaque, japanese macaque, green monkey) as our input sequences. Because PAML requires a Newick tree as an input, the MEGAX maximum likelihood algorithm was used to generate a Newick tree with the above described *SUPYN* sequences (*74*–*76*). *Baseml* was run using models 3-7 (F84, HKY85, T92, TN93, REV). As shown in **fig S16**, both the consensus- and maximum likelihood reconstructions were identical for the OWM *SUPYN* sequences. The consensus-based hominoid sequence reconstruction differed from our maximum likelihood-based reconstruction by two amino acids. These two positions are unlikely to affect the function of the resulting protein because these sites are identical to siamang *SUPYN*, which restricts RD114env-mediated infection **(Fig 4a, b)**.

## Supporting information

raw data files for SUPYN selection analysis

reconstructed ancestral SUPYN sequences

sequences of inserts cloned into pHCMV expression vector

envORF genome annotations

## Code availability

Code used to process and plot the high throughput datasets are available at https://github.com/Manu-1512/Envelopes_Suppressyn

## Acknowledgements

This work was funded by Cornell University and by the National Institutes of Health to CF (R01 GM112972, R35 GM122550). J.L.G.P’s lab acknowledge support from CICE-FEDER-P12-CTS-2256, The Wellcome Trust-University of Edinburgh Institutional Strategic Support Fund (ISFF2), the European Research Council (ERC-Consolidator ERC-STG-2012-309433), an International Early Career Scientist grant from the Howard Hughes Medical Institute (IECS-55007420), and from a private donation to the lab from Ms Francisca Serrano (*Trading y Bolsa para Torpes*, Granada, Spain). We thank Dr. Vicente Planelles for sharing reporter virus plasmids and training; Dr. Welkin Johnson for sharing his SMRVenv expression plasmid; Dr. Lars Aagaard for sharing shRNA transduction constructs; Dr. John Lis for providing lentiviral packaging construct; Dr. Amnon Koren and Rita Rebello for providing embryonic stem cell cultures and technical support respectively; Dr Lucia Carbone and Dr. Mariam Okhovat for sharing gibbon genome sequences. We thank Dr. Ray Malfavon-Borja for producing an initial list of envelope open reading frames in the human genome. We thank Maia Clare for her contribution to evolutionary sequence analyses. We thank Dr. Akiko Iwasaki for her critical reading of this manuscript. We thank members of the Feschotte lab for helpful advice and discussion throughout the project.

## Author contributions

CF conceived of this project. JAF and CF designed and developed this project. JAF designed and conducted all experiments, evolutionary sequence analyses, and analyzed all experimental data. MS performed all gene expression and regulation analyses. HBC and RAK helped perform infection assays and evolutionary sequence analyses. CBC performed placental stains. JLC and JLGP obtained, cultured, and prepared human embryos for microscopy. MBB performed embryo stains and imaging. JAF, CF and MS wrote this manuscript.

