## supplementary data for "Evolution and antiviral activity of a human protein of retroviral origin"

### Supplementary Materials

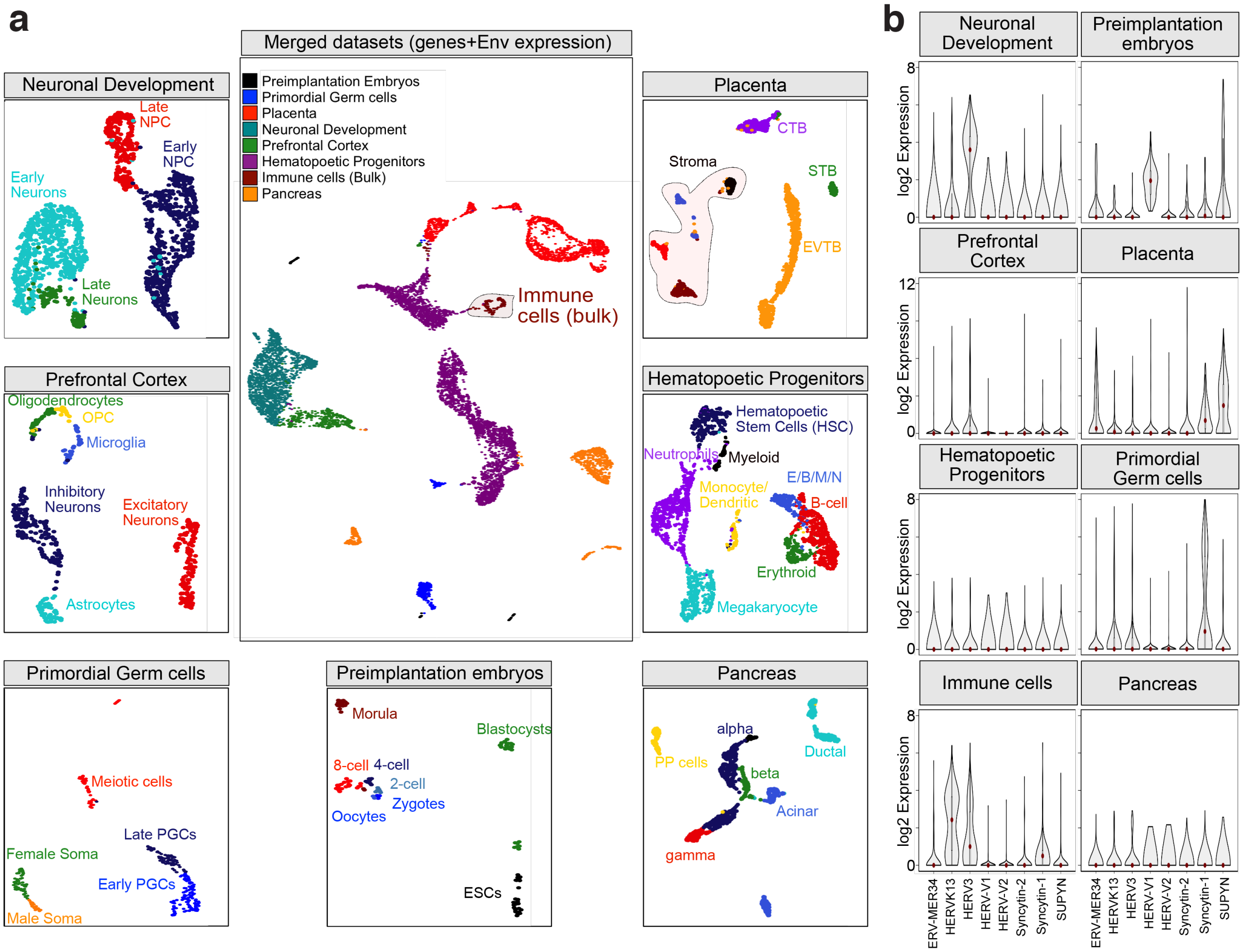


**Figure S1: Analysis of envORF expression in single cell RNA-seq taken from various human tissue sources**

**(a)** UMAP plots illustrate the Louvain clustering of seven independent sc-RNA-seq datasets corresponding to the development of human embryos and somatic tissues (see Methods). The large central UMAP plot represents our integrative analysis of seven scRNA-seq and one bulk RNA-seq datasets obtained from independent studies (Supplemental Table 1). Surrounding UMAP plots represent the combined expression of human RefSeq and envORF genes (supplemental Table 3) with cell identities labeled. **(b)** Multiple violin plots demonstrate the expression of annotated ERV envelopes in each dataset shown in **a**.

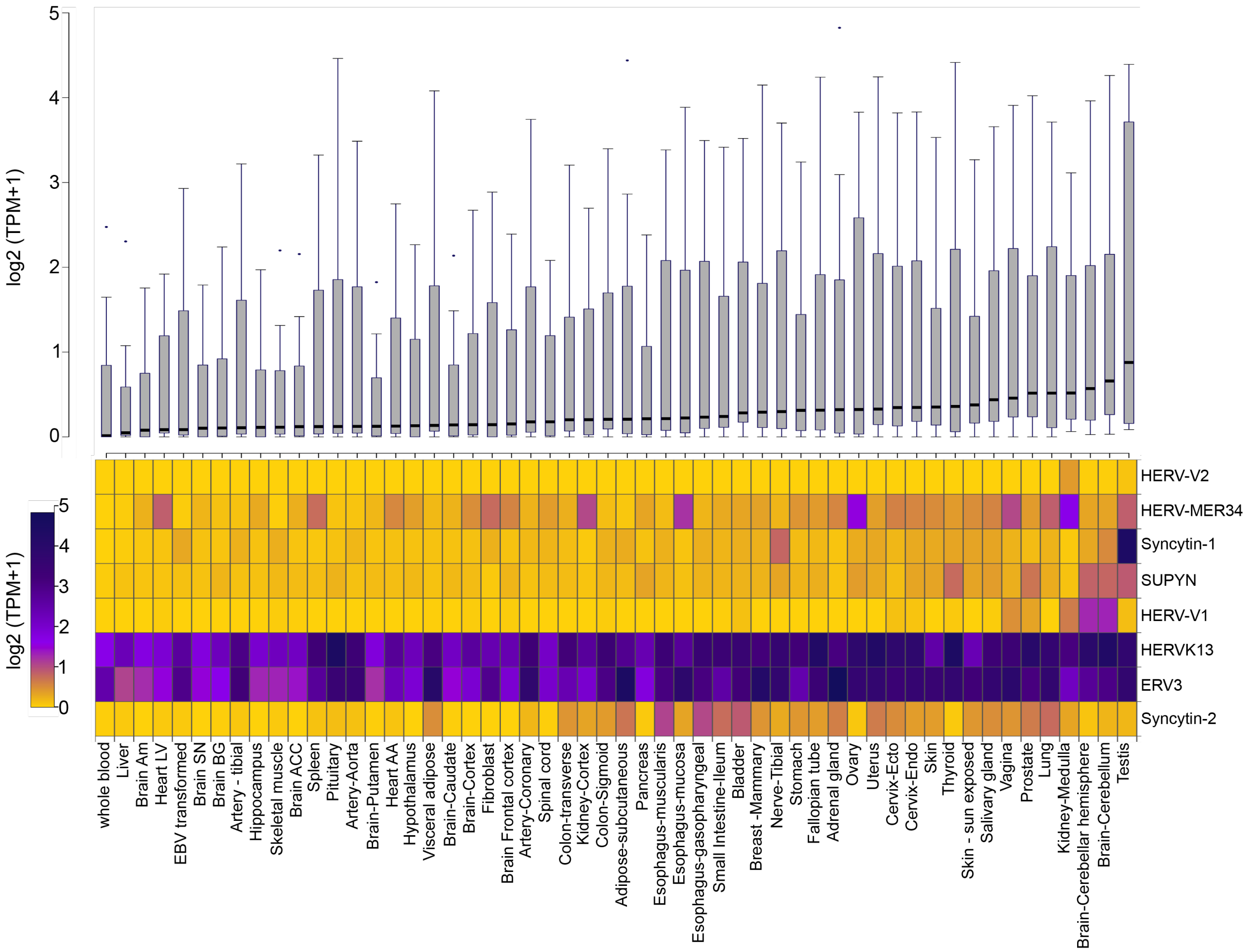


**figure S2: Annotated human endogenous retrovirus envelope expression in various human tissues.**

**(a)** The boxplot shows the transcript expression distribution of annotated ERV envelopes in tissues assayed by the GTEx project (Supplementary Table 1). **(b)** The heatmap displays the expression (log2 TPM) of individual ERV envelope genes (rows) in each tissue (columns). The color scheme ranges gradually from no expression (gold) to higher expression (midnight blue).

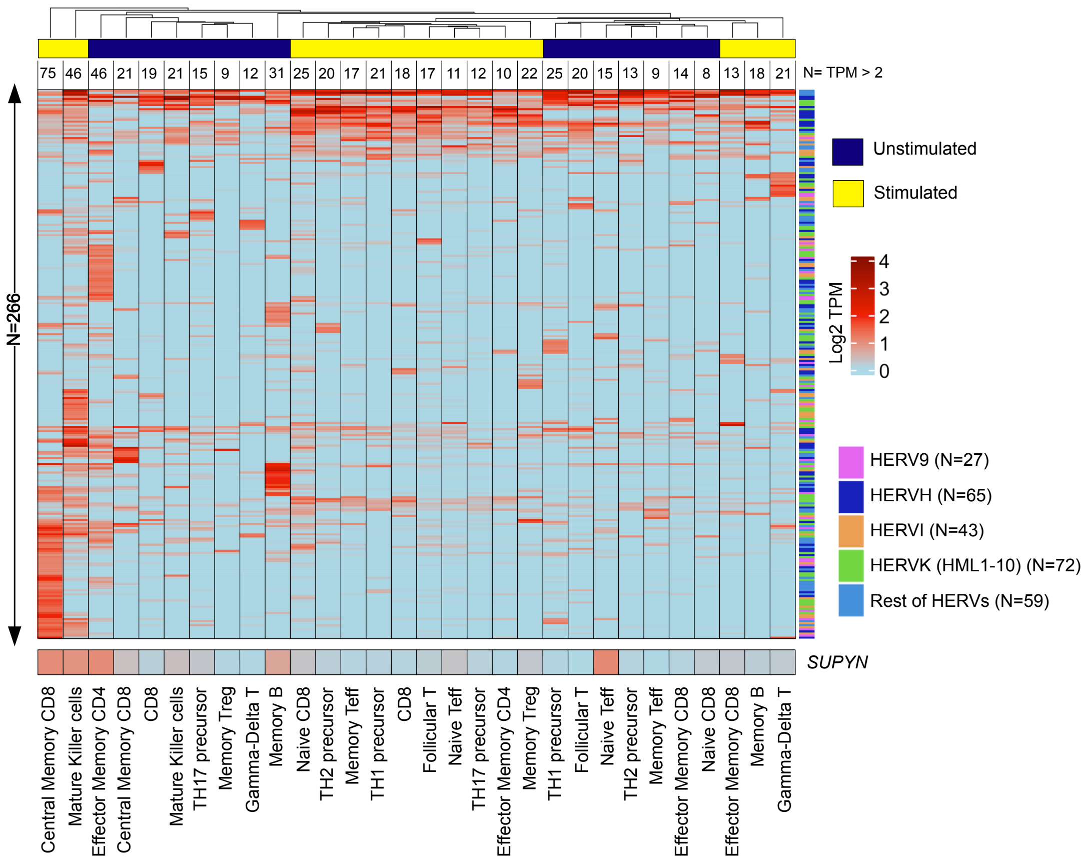


**figure S3: EnvORF expression in unstimulated and stimulated human immune cells**
Heatmap represents all envORF (N=266) with evidence of expression (log2 CPM > 1 in at least one cell type). Rows and columns represent individual *envORF* loci and cell identity respectively. Bar located above heatmap denotes unstimulated (blue) or stimulated cells (yellow) as shown in Figure 1b. Rows and columns are ordered by hierarchical clustering based on *envORF* expression.

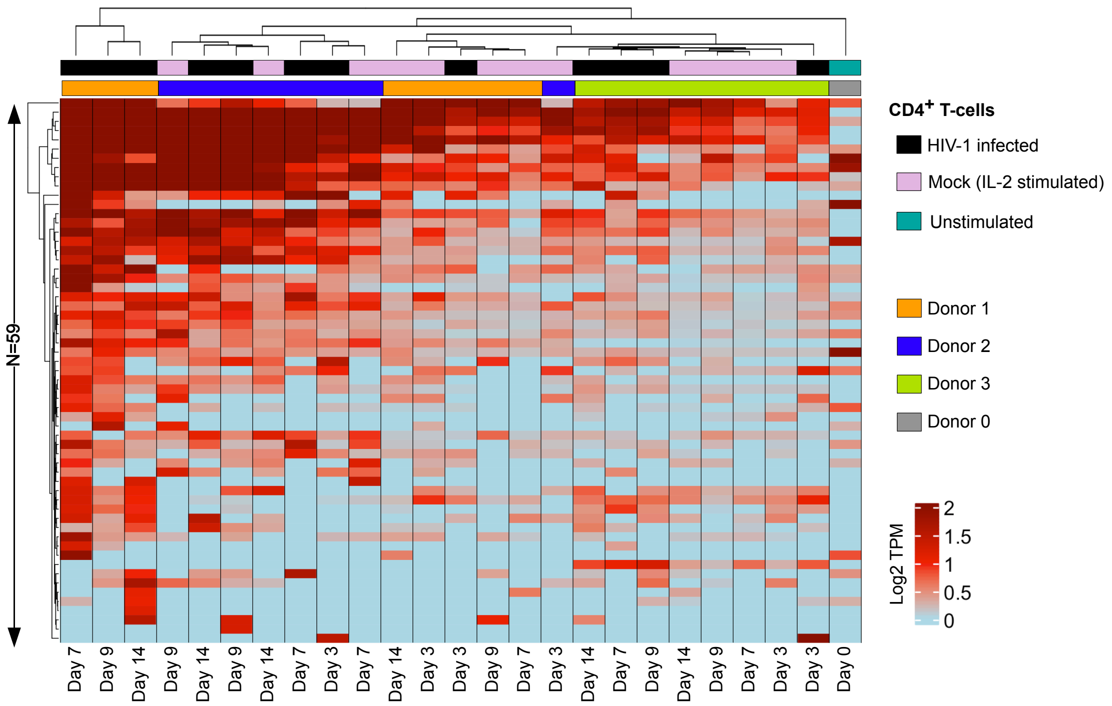


**figure S4: EnvORF expression in human T-cells in response to HIV-1 infection**

Heatmap represents all envORF (N=59) with evidence of expression (log2 CPM > 1 in at least one cell type). Rows and columns represent individual *envORF* loci and cell identity respectively. Bars located above heatmaps denote T-cell treatment (top) and donor source (bottom) as shown in Figure 1c. Rows and columns are ordered by hierarchical clustering based on *envORF* expression.

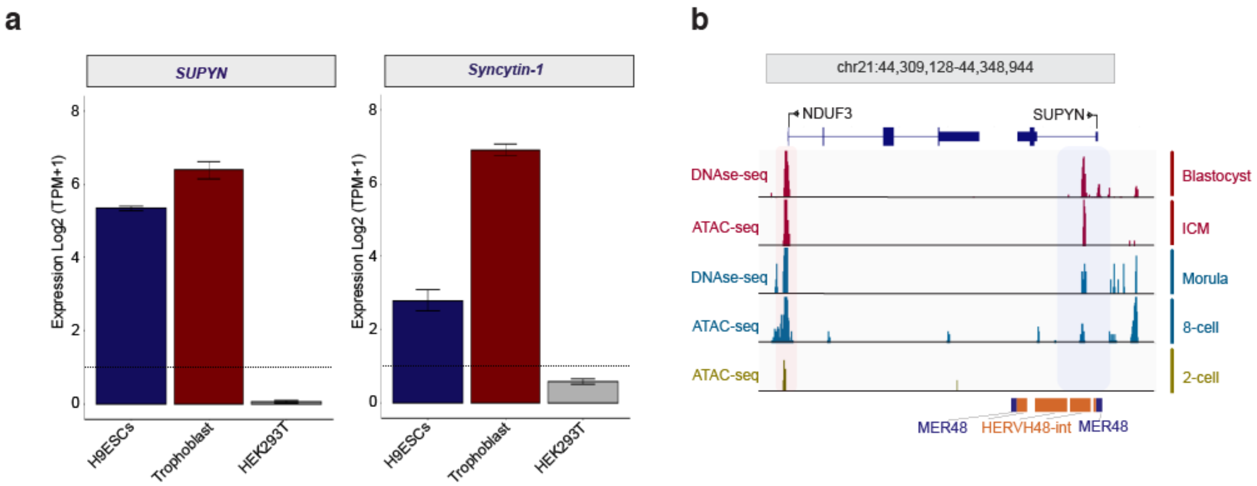


**figure S5: SUPYN is expressed in human early embryo and placenta but not in 293T cells.**
**(a)** Bar graphs show *SUPYN* and *SYN1* expression in indicated cell types. **(b)** Genome browser view showing ATAC-seq and DNAse-seq signals at the *SUPYN* locus, including upstream and downstream sequences. The framed region highlights overlapping peaks at the *SUPYN* locus.

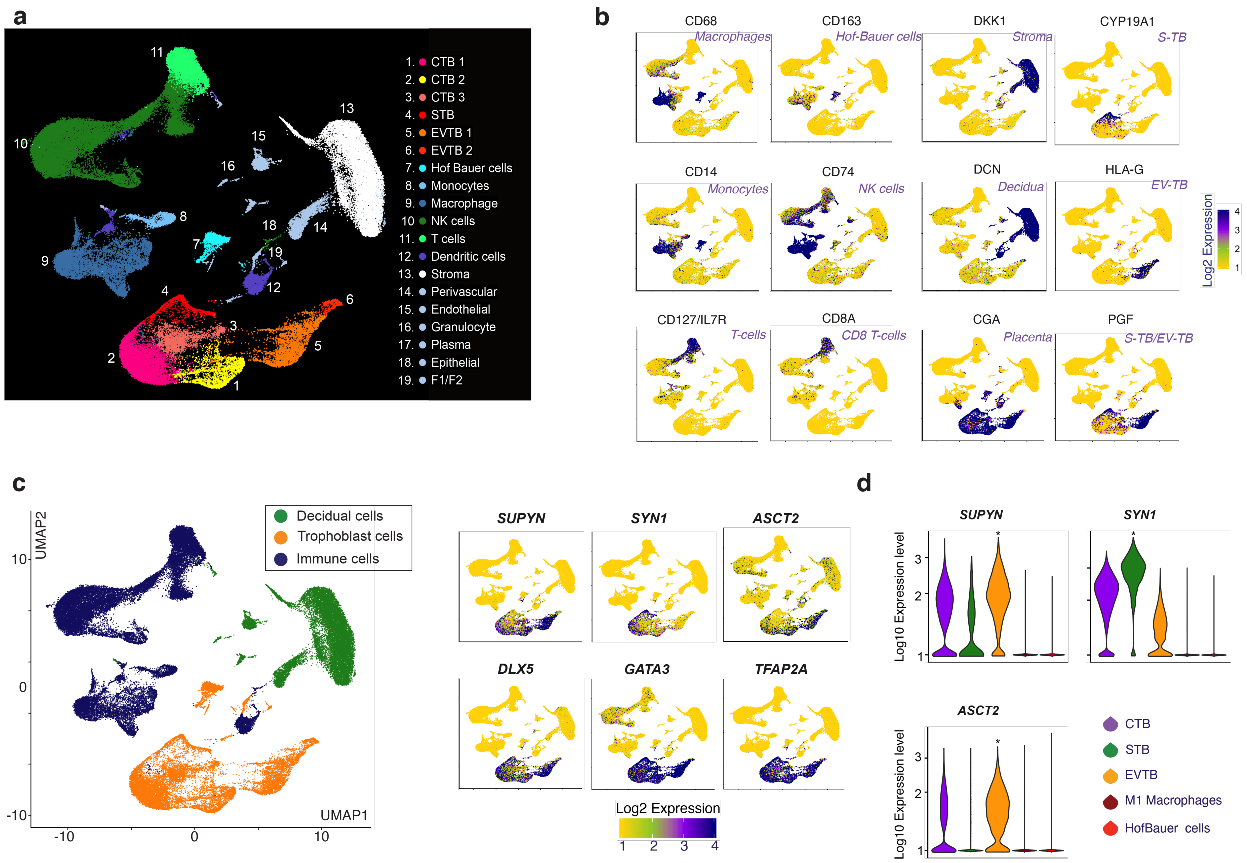


**figure S6: Defining lineage-specific SYN1, SUPYN, and ASCT2 expression from placental single-cell transcriptomics**

**(a)** UMAP plot generated from published scRNA-seq data generated from 1st trimester placental explants. Colors denote placental (pink, red, orange, and yellow) immune (blue and green) and maternal cell lineages (white and grey). **(b)** Feature plots visualize single-cell expression level of lineage-defining marker genes. **(c)** Simplified UMAP plot, shown in (a), of scRNAseq data displaying trophoblast (yellow), decidual (green) and immune (purple) cell identity. Sub-panels display single-cell-level expression of indicated genes. **(d)** Violin plots denote single-cell *SUPYN* and *ASCT2* expression in multiple placental-cell lineages.

Day 27 human blast

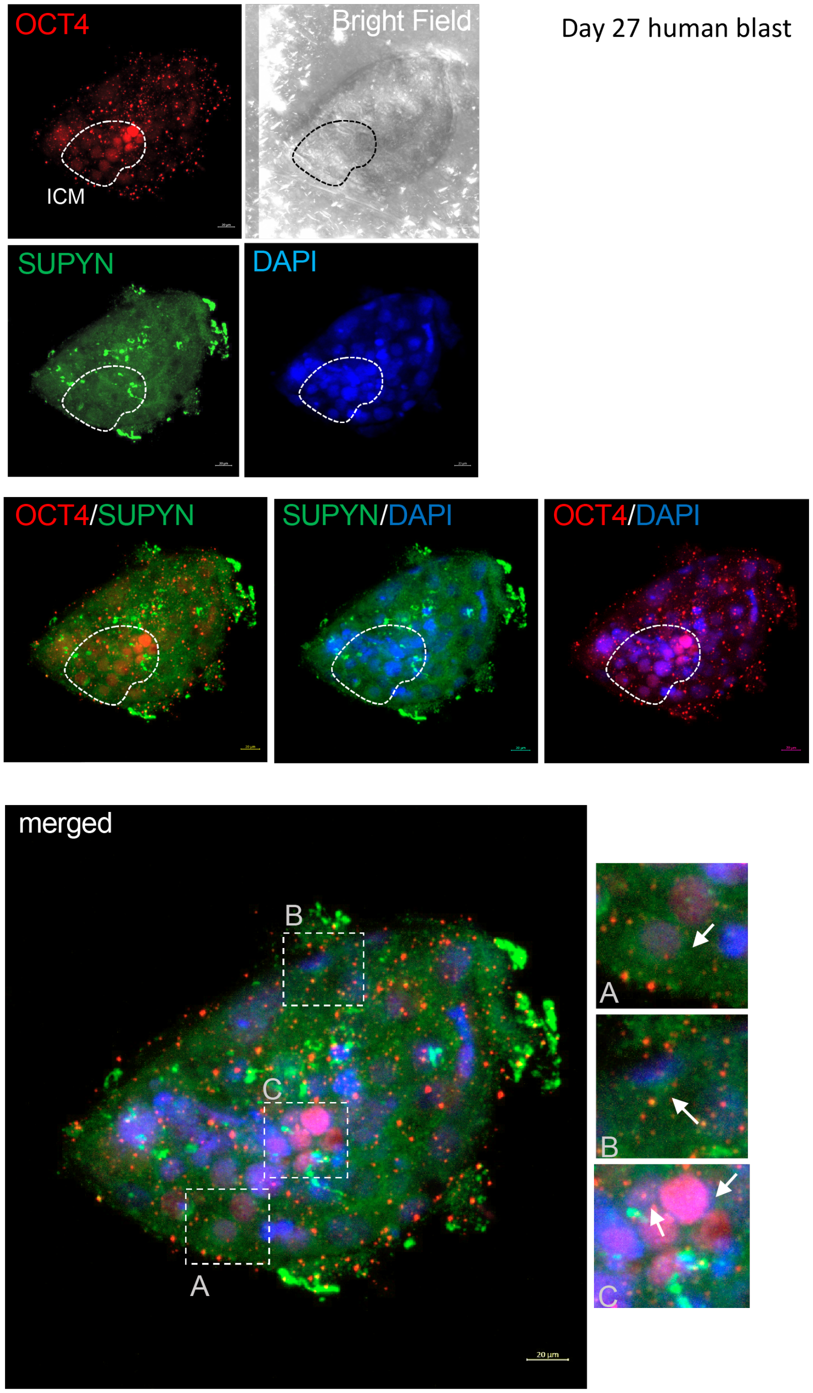


**figure S7: SUPYN and OCT4 expression in human blast stage embryos.**
Confocal microscopy of a blast stage human embryo. Embryo was immunostained for OCT4 (red) and SUPYN (green). The nucleus was labeled with DAPI (blue).

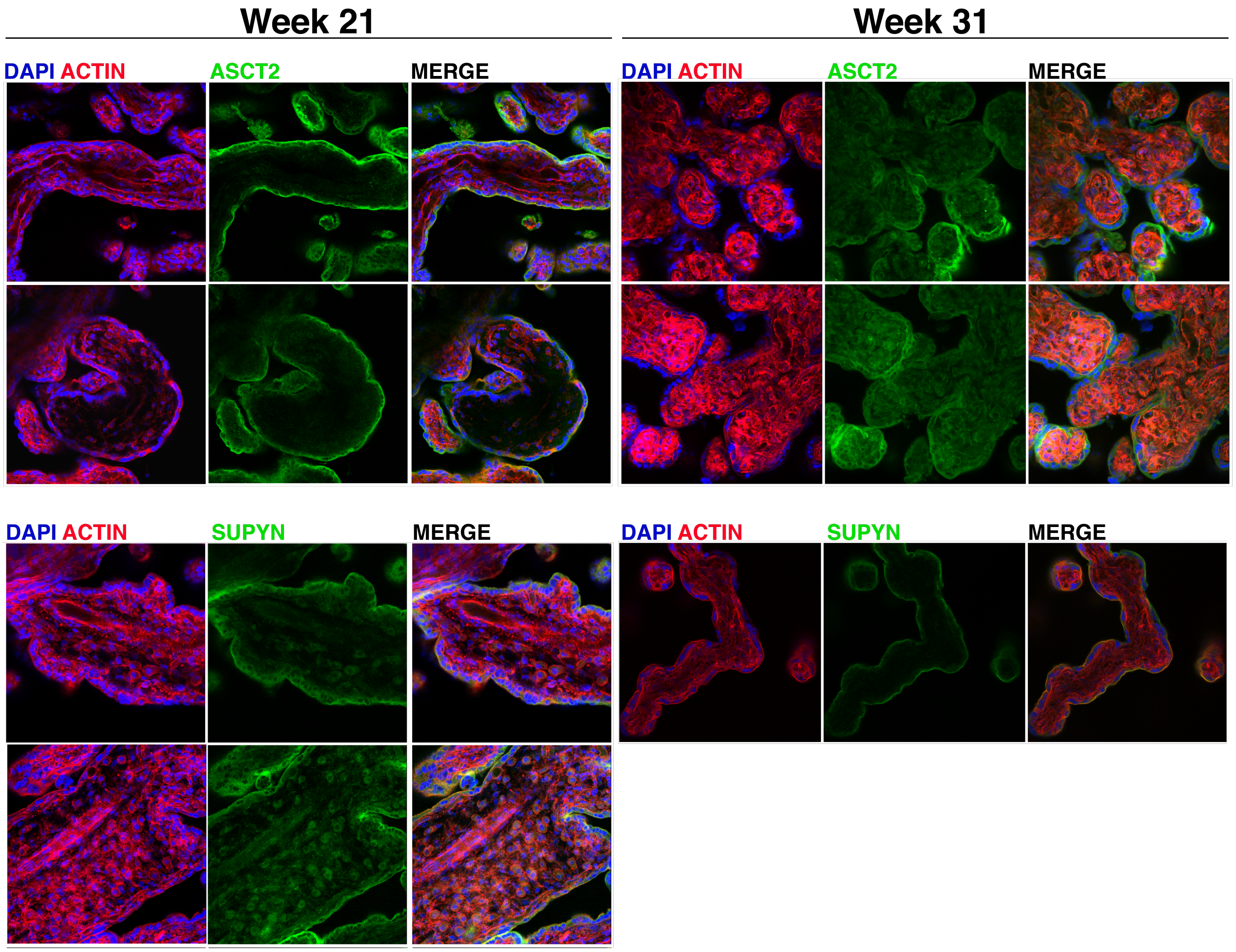


**figure S8: ASCT2 and SUPYN expression in 2nd and 3rd trimester human placenta.** Confocal microscopy of 2nd (week 21) and 3rd (week 31) trimester placental villi explants. Villi were stained for ASCT2 (green upper panels) or SUPYN (green lower panels) and Actin (red). Cell nuclei are marked with DAPI (blue).

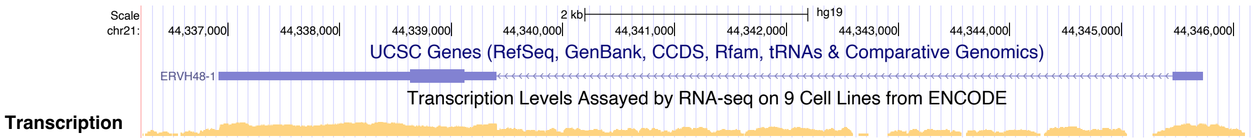


**figure S9: SUPYN expression in H1-ESCs**

Adapted UCSC genome browser shot of the SUPYN locus in the hg19 assembly UCSC Genes and ENCODE H1-ESC transcription tracks are shown. UCSC genome browser session URL: https://genome.ucsc.edu/cgi-bin/hgTracks?db=hg19&lastVirtModeType=default&lastVirtModeExtraState=&virtModeType=default&virtMode=0&nonVirtPosition=&position=chr21%3A44336232%2D44346155&hgsid=1327438861_rHJ0AIrR6S6f3iZ9r9tJbAvA7aA7

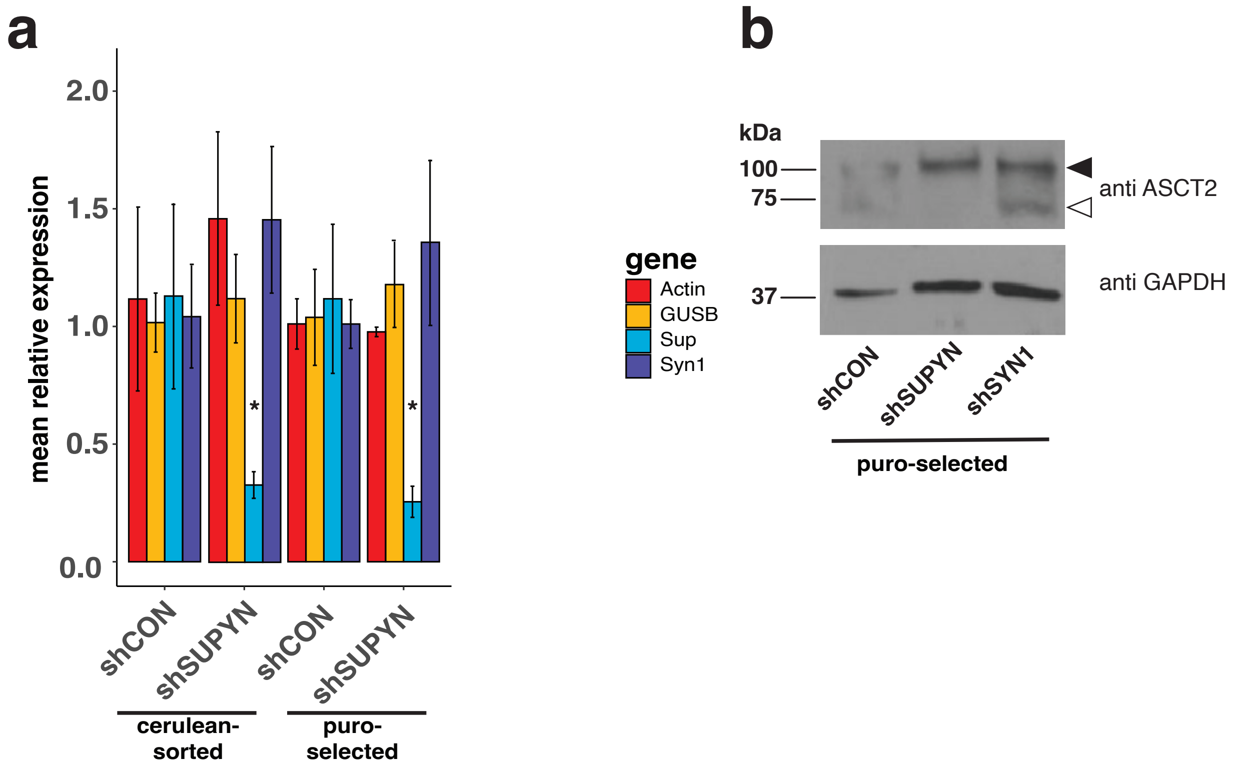


**figure S10: Characterization of shSUPYN transduced Jar cells.**
**(a)** *SUPYN* knock down was validated by qPCR. Bar plots represent mean relative gene expression normalized to shCON in cerulean-sorted and puromycin-selected cell lines respectively (n = 3). Error bars represent ± standard error mean (*$p < 0.1$; Wilcox rank sum test). **(b)** Western Blot analysis (αGAPDH, αASCT2) of shRNA-transduced Jar cell lysates. Putatively glycosylated and unglycosylated ASCT2 are marked by filled and empty arrowheads respectively.

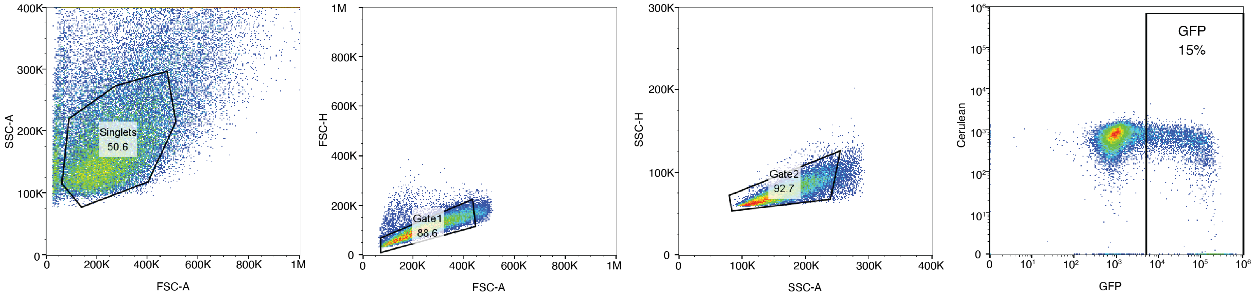


**figure S11: Flow Cytometry analysis scheme.** Representative sequential gating scheme to assess reporter virus infection rate.

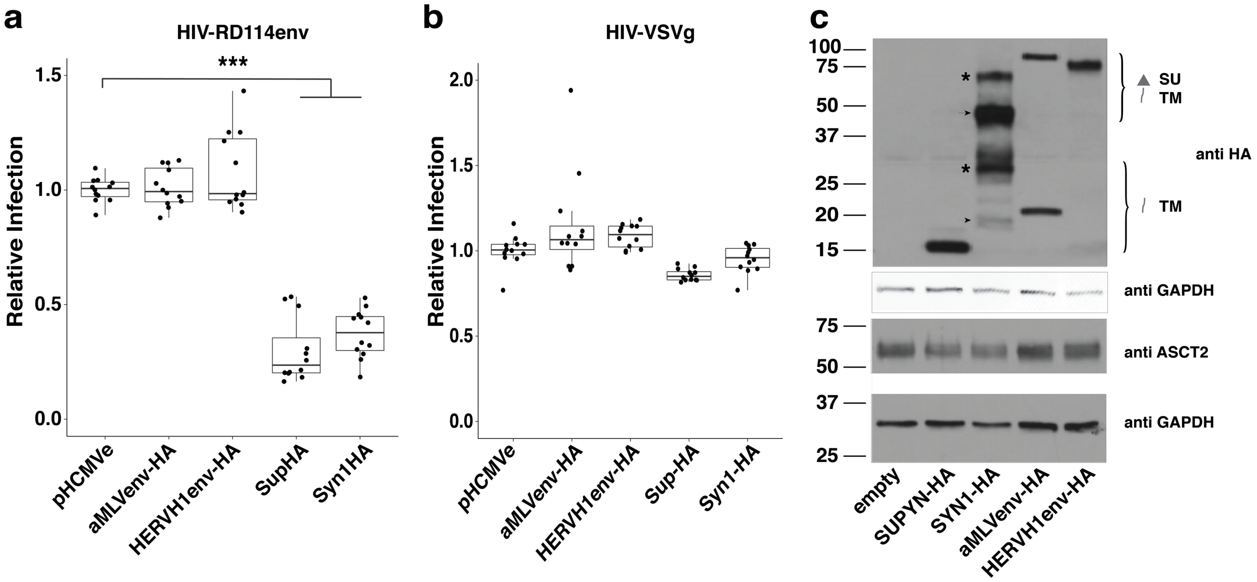


**figure S12: SUPYN expression is sufficient to specifically restrict RDR env-mediated infection**

**(a, b)** 293T cells, transfected with HA-tagged SUPYN and env constructs, were infected with HIV-RD114env **(a)** and -VSVg **(b)** respectively. Relative infection rates were determined by normalizing GFP+ counts to empty vector. (n ≥ 3 with ≥ 1 technical replicate; ***adj. $p < 0.001$; Tukey HSD). **(c)** Western Blot analysis (αHA, αGAPDH, αASCT2) of 293T cell lysates following transfection with indicated constructs. Arrowheads and asterisks denote unglycosylated and glycosylated protein fragments respectively.

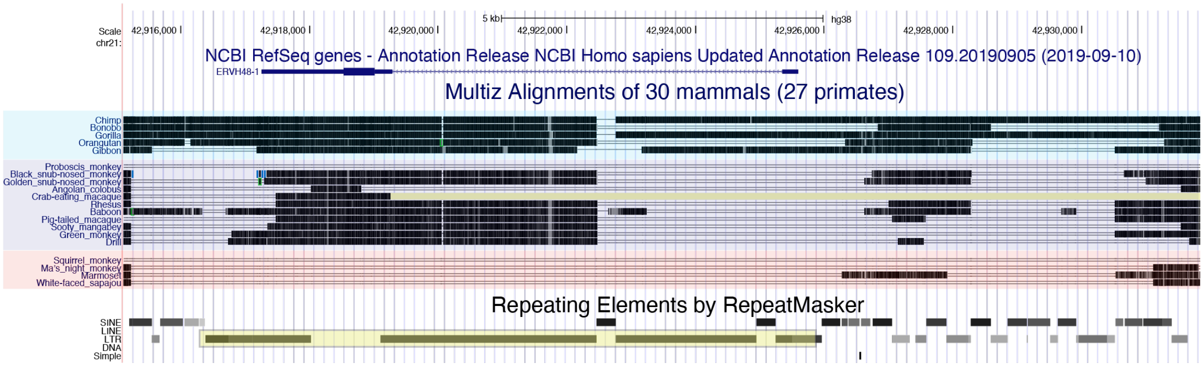


**figure S13: SUPYN locus conservation in primates.**
UCSC genome Browser snapshot of *SUPYN*-coding locus with surrounding sequence. Modified NCBI RefSeq gene, simian whole genome alignment (from Multiz 30-species track), and RepeatMasker repetitive element tracks are shown. The SUPYN-coding HERVH48 provirus is highlighted by the yellow box.

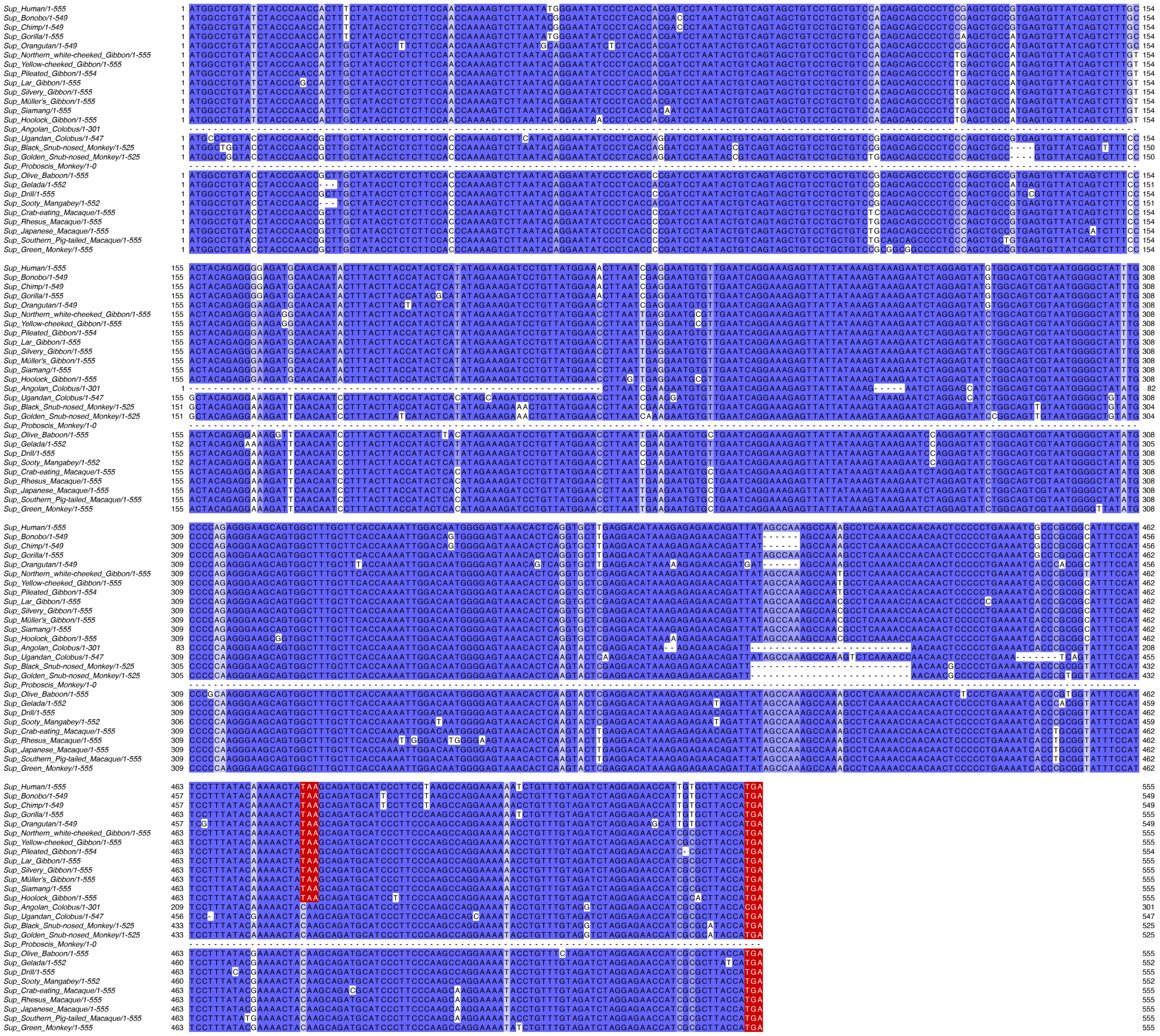

**figure S14: Nucleic acid sequence alignment of primate Suppressyn orthologs.** Suppressyn encoding nucleotide sequences are shaded blue based on a minimum sequence identity threshold of 45% (light), 75% (medium) and 80% (dark). Conserved ape-specific and ancestral stop codons are highlighted in red.

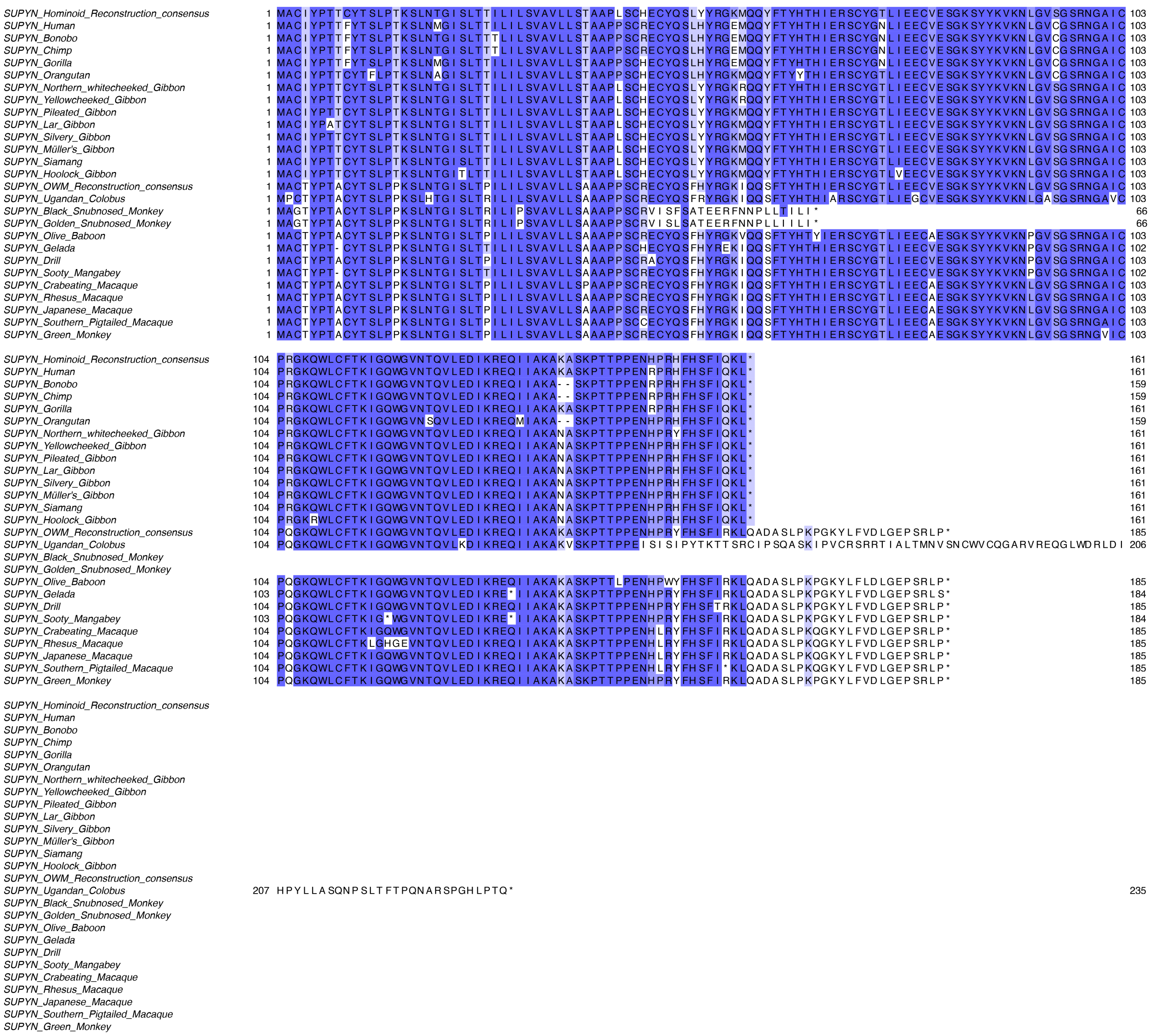

**figure S15: Amino Acid sequence alignment of primate SUPYN orthologs.** Primate and ancestral SUPYN peptide sequences are shown. Sequences are shaded blue based on a minimum sequence identity threshold of 45% (light), 75% (medium) and 80% (dark) Ancestral SUPYN sequences are based on our consensus-based sequence reconstruction (see Methods).

```
SUPYN_Hominoid_Reconstruction_consensus       1  MACIYPTTCYTSLPTKSLNTGISLTTILILSVAVLLSTAAPLSCHECYQSLYYRGKMQQYFTYHTHIERSCYGTLIEECVESGKSYYKVKNLGVSGSR  98
SUPYN_Hominoid_Reconstruction_PAML_Model3     1  MACIYPTTCYTSLPTKSLNTGISLTTILILSVAVLLSTAAPPSCHECYQSLHYRGKMQQYFTYHTHIERSCYGTLIEECVESGKSYYKVKNLGVSGSR  98
SUPYN_Hominoid_Reconstruction_PAML_Model4     1  MACIYPTTCYTSLPTKSLNTGISLTTILILSVAVLLSTAAPPSCHECYQSLHYRGKMQQYFTYHTHIERSCYGTLIEECVESGKSYYKVKNLGVSGSR  98
SUPYN_Hominoid_Reconstruction_PAML_Model5     1  MACIYPTTCYTSLPTKSLNTGISLTTILILSVAVLLSTAAPPSCHECYQSLHYRGKMQQYFTYHTHIERSCYGTLIEECVESGKSYYKVKNLGVSGSR  98
SUPYN_Hominoid_Reconstruction_PAML_Model6     1  MACIYPTTCYTSLPTKSLNTGISLTTILILSVAVLLSTAAPPSCHECYQSLHYRGKMQQYFTYHTHIERSCYGTLIEECVESGKSYYKVKNLGVSGSR  98
SUPYN_Hominoid_Reconstruction_PAML_Model7     1  MACIYPTTCYTSLPTKSLNTGISLTTILILSVAVLLSTAAPPSCHECYQSLHYRGKMQQYFTYHTHIERSCYGTLIEECVESGKSYYKVKNLGVSGSR  98
SUPYN_OWM_Reconstruction_consensus            1  MACTYPTACYTSLPPKSLNTGISLTPILILSVAVLLSAAAPPSCRECYQSFHYRGKIQQSFTYHTHIERSCYGTLIEECVESGKSYYKVKNLGVSGSR  98
SUPYN_OWM_Reconstruction_PAML_Model3          1  MACTYPTACYTSLPPKSLNTGISLTPILILSVAVLLSAAAPPSCRECYQSFHYRGKIQQSFTYHTHIERSCYGTLIEECVESGKSYYKVKNLGVSGSR  98
SUPYN_OWM_Reconstruction_PAML_Model4          1  MACTYPTACYTSLPPKSLNTGISLTPILILSVAVLLSAAAPPSCRECYQSFHYRGKIQQSFTYHTHIERSCYGTLIEECVESGKSYYKVKNLGVSGSR  98
SUPYN_OWM_Reconstruction_PAML_Model5          1  MACTYPTACYTSLPPKSLNTGISLTPILILSVAVLLSAAAPPSCRECYQSFHYRGKIQQSFTYHTHIERSCYGTLIEECVESGKSYYKVKNLGVSGSR  98
SUPYN_OWM_Reconstruction_PAML_Model6          1  MACTYPTACYTSLPPKSLNTGISLTPILILSVAVLLSAAAPPSCRECYQSFHYRGKIQQSFTYHTHIERSCYGTLIEECVESGKSYYKVKNLGVSGSR  98
SUPYN_OWM_Reconstruction_PAML_Model7          1  MACTYPTACYTSLPPKSLNTGISLTPILILSVAVLLSAAAPPSCRECYQSFHYRGKIQQSFTYHTHIERSCYGTLIEECVESGKSYYKVKNLGVSGSR  98

SUPYN_Hominoid_Reconstruction_consensus      99  NGAICPRGKQWLCFTKIGQWGVNTQVLEDIKREQIIAKAKASKPTTPPENHPRHFHSFIQKL*ADASLPKPGKNLFVDLGEPSRLP*  185
SUPYN_Hominoid_Reconstruction_PAML_Model3    99  NGAICPRGKQWLCFTKIGQWGVNTQVLEDIKREQIIAKAKASKPTTPPENHPRHFHSFIQKLQADASLPKPGKYLFVDLGEPSRLP-  184
SUPYN_Hominoid_Reconstruction_PAML_Model4    99  NGAICPRGKQWLCFTKIGQWGVNTQVLEDIKREQIIAKAKASKPTTPPENHPRHFHSFIQKLQADASLPKPGKYLFVDLGEPSRLP-  184
SUPYN_Hominoid_Reconstruction_PAML_Model5    99  NGAICPRGKQWLCFTKIGQWGVNTQVLEDIKREQIIAKAKASKPTTPPENHPRHFHSFIQKLQADASLPKPGKYLFVDLGEPSRLP-  184
SUPYN_Hominoid_Reconstruction_PAML_Model     99  NGAICPRGKQWLCFTKIGQWGVNTQVLEDIKREQIIAKAKASKPTTPPENHPRHFHSFIQKLQADASLPKPGKYLFVDLGEPSRLP-  184
SUPYN_Hominoid_Reconstruction_PAML_Model7    99  NGAICPRGKQWLCFTKIGQWGVNTQVLEDIKREQIIAKAKASKPTTPPENHPRHFHSFIQKLQADASLPKPGKYLFVDLGEPSRLP-  184
SUPYN_OWM_Reconstruction_consensus           99  NGAICPQGKQWLCFTKIGQWGVNTQVLEDIKREQIIAKAKASKPTTPPENHPRYFHSFIRKLQADASLPKPGKYLFVDLGEPSRLP*  185
SUPYN_OWM_Reconstruction_PAML_Model3         99  NGAICPQGKQWLCFTKIGQWGVNTQVLEDIKREQIIAKAKASKPTTPPENHPRYFHSFIRKLQADASLPKPGKYLFVDLGEPSRLP-  184
SUPYN_OWM_Reconstruction_PAML_Model4         99  NGAICPQGKQWLCFTKIGQWGVNTQVLEDIKREQIIAKAKASKPTTPPENHPRYFHSFIRKLQADASLPKPGKYLFVDLGEPSRLP-  184
SUPYN_OWM_Reconstruction_PAML_Model5         99  NGAICPQGKQWLCFTKIGQWGVNTQVLEDIKREQIIAKAKASKPTTPPENHPRYFHSFIRKLQADASLPKPGKYLFVDLGEPSRLP-  184
SUPYN_OWM_Reconstruction_PAML_Model6         99  NGAICPQGKQWLCFTKIGQWGVNTQVLEDIKREQIIAKAKASKPTTPPENHPRYFHSFIRKLQADASLPKPGKYLFVDLGEPSRLP-  184
SUPYN_OWM_Reconstruction_PAML_Model7         99  NGAICPQGKQWLCFTKIGQWGVNTQVLEDIKREQIIAKAKASKPTTPPENHPRYFHSFIRKLQADASLPKPGKYLFVDLGEPSRLP-  184
```

**Figure S16: Amino Acid sequence alignment of ancestral SUPYN sequences.** Peptide sequences of consensus and maximum likelihood based SUPYN sequence reconstructions (see methods) are aligned.

**Tables**

**Table S1: External data sources**

| Description | Author | Year | PMID | Ref. | Dataset | Tissue source | Fig. |
|---|---|---|---|---|---|---|---|
| scRNA-seq | Yan et al. | 2013 | 23934149 | 31 | GSE36552 | embryo | 1a, 2a, S1 |
| | Vento-Tormo et al. | 2018 | 30429548 | 35 | E-MTAB-6701 | placenta | 2d-f, S5 |
| | Liu et al. | 2018 | 30042384 | 33 | GSE89497 | placenta | 1a, 2f, S1 |
| | Pavlicev et al. | 2017 | 28174237 | 34 | GSE87726 | placenta | 2f |
| | Close | 2017 | 28279351 | 36 | GSE93593 | Neuronal differentiation | 1a, S1 |
| | Guo | 2015 | 26046443 | 32 | GSE63818 | PGC | 1a, S1 |
| | Velten | 2017 | 28319093 | 37 | GSE75478 | HSC | 1a, S1 |
| | Enge | 2017 | 28965763 | 38 | GSE81547 | Pancreas | 1a, S1 |
| | Darmanis | 2015 | 26060301 | 39 | GSE67835 | PFC | 1a, S1 |
| Bulk RNA-seq | GTEx Consortium | 2017 | 29022597 | 45 | phs000424.v6.p1 | Human Tissues | S2 |
| | Calderon et al | 2019 | 31570894 | 49 | GSE118165 | Immune cells | 1b, S1, S3 |
| | Shytaj et al | 2020 | 32157726 | 50 | GSE12746 | CD4 T-cells | 1c, S3 |
| | Zadora et al. | 2017 | 28904069 | 46 | available via approval | placenta | 2f |
| | D Antonio et al. | 2017 | 28874753 | 47 | E-MTAB-5714 | 293T | S4a |
| | Chung et al. | 2018 | 29395325 | 48 | GSE99249 | 293T | S4a |
| ChIPseq | Barakat et al. | 2018 | 30033119 | 52 | GSE99631 | hESC | 2b |
| | Krendl et al. | 2017 | 29078328 | 55 | GSE105258 | hESC | 2b, S4a-b |

| | | | | | | | |
|---|---|---|---|---|---|---|---|
| | Krendl et al. | 2017 | 29078328 | 55 | GSE105081 | hESC | 2b, S4a-b |
| | Tsankov et al. | 2015 | 25693565 | 51 | GSE61475 | hESC | 2b, S4d |
| | Dunn-Fletcher et al. | 2018 | 30231016 | 54 | GSE118289 | placenta | 1c |
| | Kwak et al. | 2019 | 31294776 | 53 | GSE127288 | placenta | 1c |
| DNAse-seq | Gao et al. | 2018 | 29526463 | 56 | GSA:CRA000297 | embryo | S1c |
| ATAC-seq | Wu et al. | 2018 | 29720659 | 57 | GSE101571 | embryo | S1c |

**Table S2: Primer list**

| InFusion cloning primers | | |
|---|---|---|
| **ID** | **sequence** | **description** |
| JF023 | TTTTGGCAAAGAATTCATGGCCTGTATCTACCCA | pHCMV human suppressyn fwd |
| JF024 | CCTGAGGAGTGAATTCTTATAGTTTTTGTATAAAGGAATGG | pHCMV human suppressyn rev |
| JF025 | TTTTGGCAAAGAATTCATGGCCTGTATCTACCCAACC | pHCMV human suppressyn-HA fwd |
| JF026 | CCTGAGGAGTGAATTCTTAAGCGTAATCTGGAACATCG | pHCMV human suppressyn-HA rev |
| JF046 | TTTTGGCAAAGAATTCATGGCGCGTTCAACGCTCT | pHCMV amphoMLVenv fwd |
| JF047 | CCTGAGGAGTGAATTCTCATGGCTCGTACTCTATGGGT | pHCMV amphoMLVenv rev |
| JF081 | GCATAGTAGTCTCATTGCTACCACA | HERVH-1 (H62) provirus fwd |
| JF082 | CATGACTCGGATCAGGGGAC | HERVH-1 (H62) provirus rev |
| JF091 | ATCATTTTGGCAAAGaattCATGATCTTTGCTGGCAAGGCACC | pHCMV HERVH-1env fwd |
| JF093 | CAGCCTGCACCTGAGGAGTGaattCctaagcgtaatctggaacatcgtatgggtaAGCTGAAGGGAGGTCTTGTGGTAAG | pHCMV HERVH-1env-HA rev |
| JF130 | GGCGCCTAAGCTGGTATTCTTAACTATGTTGCTCC | pHIV7 eGFP downstream sequence fwd |
| JF131 | CGCAGAGCCGGCAGCAGGCCGCGGGAAGGAAGGTCCGCTGGATTG | pHIV7 eGFP downstream sequence rev |
| JF135b | tttggggatcctgagcggccgcatccaaggtcgggcagga | U6 shRNA cloning fwd (pHIV-shSup cloning) |
| JF136 | ccaaccactttctatactacacaaatatagaaagtggttgggtagatacactttcgtcctttccacaagatatataaagccaagaa | U6-shSup rev (pHIV-shSup cloning) |

| | | |
|---|---|---|
| JF138 | CTACTTCCTTTCGATACTACACAAATATCGAAAG GAAGTAGAAGTCCGGGCTTTCGTCCTTTCCACAA GATATATAAAGCCAAGAA | U6-shC rev (pHIV-shC cloning) |
| JF145 | gtagtatagaaagtggttgggtagatacact tttttgtaccgagctcggatccactagagatgg | shSup-CMV promoter fwd (pHIV7-shSup cloning) |
| JF147 | GTAGTATCGAAAGGAAGTAGAAGTCCGGGCT tttttgtaccgagctcggatccactagagatgg | shC-CMV promoter fwd (pHIV7-shC cloning) |
| JF152b | TGAACCGTCAGATCCGCTAGCATGGTGAGCAAGG GCGAGG | cerulean fwd (pHIV7-shRNA-cerulean cloning) |
| JF153b | AGAATACCAGttACTTGTACAGCTCGTCCATGCC | cerulean rev (pHIV7-shRNA-cerulean cloning) |
| JF154b | CTGTACAAGTAACTGGTATTCTTAACTATGTTGC TCC | cerulean-WPRE fwd (pHIV7-shRNA-cerulean cloning) |
| JF155 | TGAACCGTCAGATCCGCTAGCATGGCCACCGAGT ACAAG | puroR fwd (pHIV7-shRNA-puroR cloning) |
| JF156 | AGAATACCAGTTATTTGTAACCATTATAAGCTGC | puroR rev (pHIV7-shRNA-puroR cloning) |
| JF157 | TTACAAATAACTGGTATTCTTAACTATGTTGCTC C | puroR-WPRE fwd (pHIV7-shRNA-puroR cloning) |
| JF165 | cggtggccatGctagcggatctgacggttc | pHIV7-CMV promoter rev (pHIV-shRNA puroR cloning) |
| JF166 | tgctcaccatgctagcggatctgacggttc | pHIV7-CMV promoter rev (pHIV-shRNA Cerulean cloning) |
| RAK010 | TTTTGGCAAAGAATTCATGCTCTGCATCCTCATC CTCCT | pHCMV SMRVenv cloning primer fwd |
| RAK011 | CCTGAGGAGTGAATTCCTACAGTCTGCCATATTC TAGGTCACGA | pHCMV SMRVenv cloning primer rev |
| **qPCR primers** | | |
| JF042 | GCAATTATTCCCCATGAACG | 18S fwd |
| JF043 | GGCCTCACTAAACCATCCAA | 18S rev |
| JF108 | CTGTGCATGCACATCGGTCACTG | Suppressyn fwd |
| JF109 | GAGAAATTGGCCCAGACAAACACT | Suppressyn rev |
| JF112 | ACCACGAACGGACATCCAAAG | Syncytin-1 fwd |
| JF113 | GCCACTTTAACCGCAGTTGG | Syncytin-1 rev |
| Act-F | CGACAGGATGCAGAAGGAG | Actin fwd |
| Act-R | GTACTTGCGCTCAGGAGGAG | Actin rev |
| GUSB-F | AGAGTGGTGCTGAGGATTGG | GUSB fwd |

| | | |
|---|---|---|
| GUSB-R | CCCTCATGCTCTAGCGTGTC | GUSB rev |
